# The cardiac TRPA1 channel drives calcium-mediated mechano-arrhythmogenesis

**DOI:** 10.1101/2020.10.01.321638

**Authors:** Breanne A. Cameron, Matthew R. Stoyek, Jessi J. Bak, Michael S. Connolly, Emma A. DeLong, Joachim Greiner, Rémi Peyronnet, Peter Kohl, T Alexander Quinn

## Abstract

Maintenance of cardiac function involves a regulatory loop in which electrical excitation causes the heart to contract through excitation-contraction coupling (ECC),^1^ and the mechanical state of the heart directly affects its electrical activity through mechano-electric coupling (MEC).^2^ However, in pathological states such as acute ischaemia that alter early or late electro-mechanical coordination (*i.e.*, disturbances in ECC or repolarisation-relaxation coupling, RRC), MEC may contribute to the initiation and / or sustenance of arrhythmias (mechano-arrhythmogenesis).^3^ The molecular identity of specific factor(s) underlying mechano-arrhythmogenesis in acute ischaemia, however, remain undefined.^4^ By rapid stretch of rabbit single left ventricular cardiomyocytes, we show that upon ATP-sensitive potassium channel-induced alterations of RRC, overall vulnerability to mechano-arrhythmogenesis is increased, with mechano-sensitive^5–11^ transient receptor potential kinase ankyrin 1 (TRPA1) channels^12^ acting as the molecular driver through a Ca^2+^-mediated mechanism. Specifically, TRPA1 activation drives stretch-induced excitation and creates a substrate for self-sustained arrhythmias, which are maintained by increased cytosolic free Ca^2+^ concentration ([Ca^2+^]_i_) and spontaneous [Ca^2+^]_i_ oscillations. This TRPA1-dependent mechano-arrhythmogenesis involves microtubules, and can be prevented by block of TRPA1 or buffering of [Ca^2+^]_i_. Thus, in cardiac pathologies with disturbed RRC dynamics and / or augmented TRPA1 activity, TRPA1 may represent an anti-arrhythmic target with untapped therapeutic potential.^13–17^

## MAIN TEXT

Feedback is an essential element of biological function and fundamental to the adaptation of physiological activity to varying demands. A prime example is found in the heart, an electro-mechanical pump wherein electrical excitation causes mechanical contraction of cardiomyocytes (CM) through a feedforward mechanism involving triggered release of calcium (Ca^2+^) from intracellular stores, known as excitation-contraction coupling.^1^ Feedback in the heart involves effects of the mechanical state of CM on their electrical activity, by a process termed mechano-electric coupling.^2^ While this feedback is important for maintaining and fine-tuning normal cardiac function, mechano-electric coupling can also contribute to arrhythmogenesis through mechanically-induced electrophysiological alterations and responses (mechano-arrhythmogenesis).^18^

In physiological conditions, the heart is protected from mechano-arrhythmogenesis at all phases of the cardiac cycle, in part through the tight coordination of normal cardiac electro-mechanical activity and transient electrical refractoriness following initiation of the action potential (AP). In disease, disruption of electro-mechanical coordination, haemodynamic overload (either preload or afterload), and / or altered myocardial mechanical properties can elicit arrhythmogenic mechano-electric coupling effects.^3^ While the influence of mechano-electric coupling on cardiac electrophysiology in health and disease is well established,^2^ the molecular identity of specific driver(s) underlying mechano-arrhythmogenesis is still being explored.^4^

Recent results from rabbit isolated heart studies indicate that ventricular mechano-arrhythmogenesis in acute regional ischaemia (as occurs with obstruction of a coronary artery) involves a mechano-sensitive Ca^2+^-mediated mechanism.^20^ The arrhythmogenic role of Ca^2+^ may be facilitated by a pathological disruption of normal electro-mechanical coordination during the late phase of the AP.^20–22^ Specifically, ischaemia can induce a dissociation of the otherwise well-synchronised post-excitation recovery of membrane potential and cytosolic free Ca^2+^ concentration ([Ca^2+^]_i_). This process, which we henceforth refer to as ‘repolarisation-relaxation coupling’ (RRC), describes the period in which membrane repolarisation prevents further voltage-mediated trans-sarcolemmal Ca^2+^ influx, and existing cytosolic Ca^2+^ is sequestered into intracellular stores or extruded from the cell, thereby eliciting relaxation. With reduced oxygen availability in ischaemia, activation of ATP-sensitive potassium (K_ATP_) channels causes rapid cellular repolarisation and an associated shortening of the [Ca^2+^]_i_ transient. The resulting decrease in action potential duration (APD) is greater, however, than that in [Ca^2+^]_i_ transient duration (CaTD), resulting in a disturbance of normal RRC.^20–22^ This may be arrhythmogenic, as it results in the emergence of a potentially vulnerable period (VP_RRC_) for Ca^2+^-mediated arrhythmogenesis during late repolarisation, when [Ca^2+^]_i_ remains elevated in progressively re-excitable tissue. Such a VP_RRC_ has been shown to occur during beta-adrenergic stimulation in failing human hearts.^23^

In the present study, we utilised a cell-level approach that allows for the control of single cell mechanical load with a carbon fibre-based system^19^ to investigate the effects of disturbed RRC and the resultant VP_RRC_ on the susceptibility of rabbit left ventricular CM to mechano-arrhythmogenesis. This was combined with fluorescence imaging of trans-membrane voltage and [Ca^2+^]_i_, video-based measurement of sarcomere dynamics, and pharmacological interrogation of mechanisms suspected to be involved in triggering and / or sustaining arrhythmic activity upon cell stretch.

To mimic ischaemic changes in AP morphology, K_ATP_ channels were activated by application of pinacidil (50 μM, continuous superfusion), a well-established agonist of sulfonylurea receptor (SUR)2A / K_ir_6.2 in cardiac and skeletal muscle.^24^ To assess effects of K_ATP_ channel activation on relative post-activation recovery-times of voltage (APD at 50 % repolarisation, APD_50_) and [Ca^2+^]_i_ (CaTD at 80 % return to diastolic levels, CaTD_80_), as well as on the duration of the resultant VP_RRC,_ we used a single-excitation / dual-emission fluorescence imaging approach.^25^ The voltage-sensitive dye di-4-ANBDQPQ and the Ca^2+^-sensitive dye Fluo-5F, AM (whose high K_d_ value [∼2.3 μM] limits its buffering of near-diastolic [Ca^2+^]_i_ and potential artefactual effects on CaTD_80_) were used to simultaneously measure AP and [Ca^2+^]_i_ transients in electrically-paced rabbit left ventricular CM under carbon fibre control (Fig. 1a). Upon pinacidil exposure, both APD and CaTD decreased (Fig. 1b, c). The decrease of APD, however, was significantly greater than that of CaTD, resulting in the formation of a cellular VP_RRC_. This period spans the time during which a CM starts to become re-excitable (repolarisation past APD_50_) while [Ca^2+^]_i_ remains elevated (above CaTD_80_; Fig. 1b, d). This response mimics behaviour previously observed in whole hearts during acute ischaemia,^20, 21^ as well as with hypoxia alone.^22^ After 5 min of pinacidil exposure, there were no further significant alterations in APD, CaTD, or the VP_RRC_ (Fig. 1c, d).

**Figure 1.**
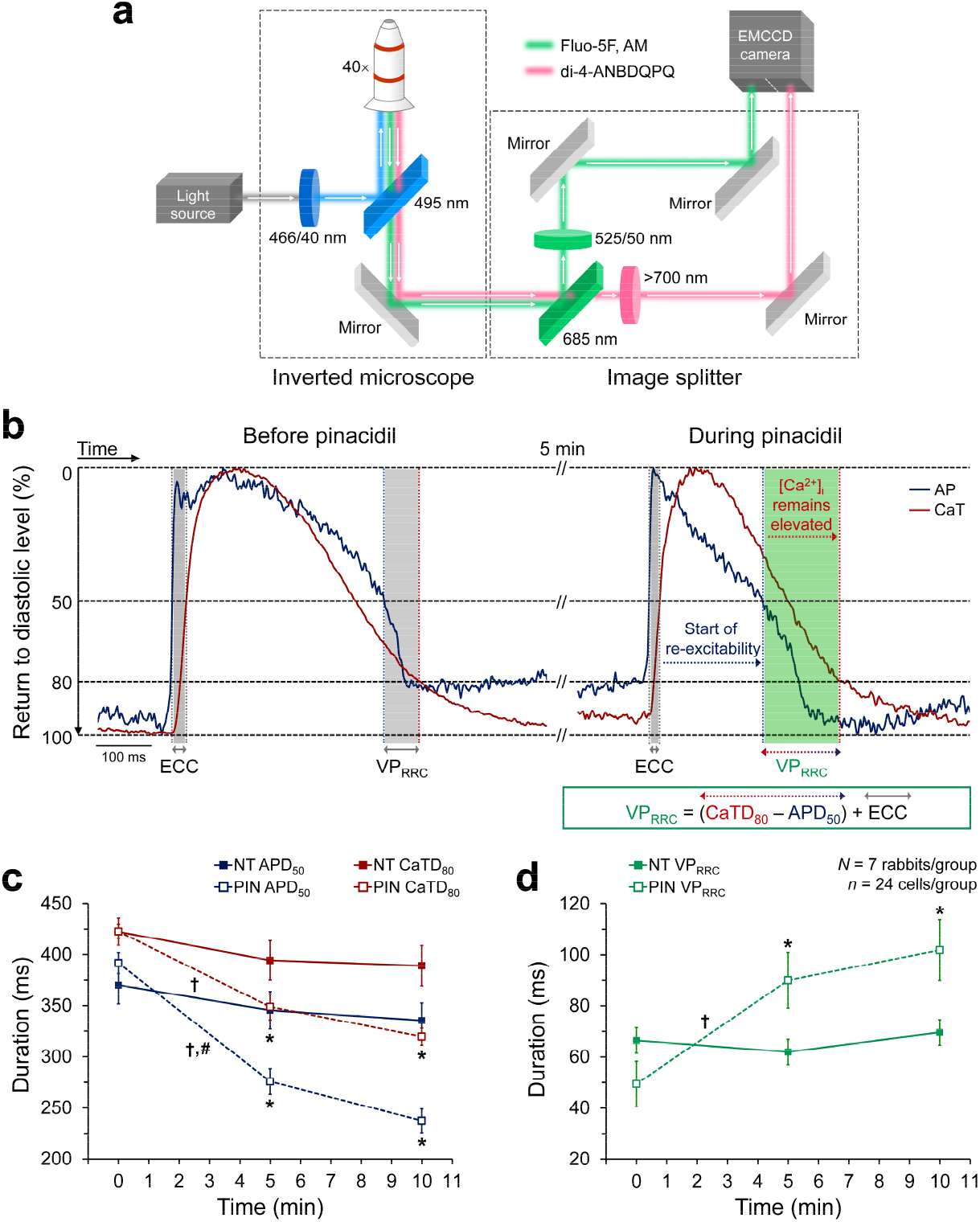
Disruption of repolarisation-relaxation coupling (RRC) in rabbit isolated left ventricular cardiomyocytes by activation of ATP-sensitive potassium channels with pinacidil. **a,** Schematic of the single-excitation/dual-emission fluorescence imaging approach, utilising a single camera-image splitter system. **b,** Representative traces of action potentials (AP, blue) and calcium transients (CaT, red), simultaneously recorded by fluorescence imaging in a single ventricular cardiomyocyte before (left) and after (right) 5 min of pinacidil exposure (50 µM, to activate ATP-sensitive potassium channels), showing RRC disruption. The associated vulnerable period (VP_RRC_; highlighted in green) is the interval during which cardiomyocytes start to become re-excitable, while cytosolic calcium concentration ([Ca^2+^]_i_) remains elevated (calculated as the difference between the time of recovery of cytosolic Ca^2+^ to 80 % [CaTD_80_, red dashed line] and membrane potential to 50 % [APD_50_, blue dashed line] of diastolic levels), plus the excitation contraction coupling time (ECC). **c,** APD_50_ (blue), CaTD_80_ (red), and d, VP_RRC_ duration (green) during 10 min of Tyrode (NT, solid) or pinacidil (PIN, dashed) superfusion. Differences assessed by one-way ANOVA with Tukey *post-hoc* tests. \**p* < 0.05 for NT compared to PIN; †*p* < 0.05 for the change within either NT or PIN over time; ^#^*p* < 0.05 for the change over time of APD_50_ compared to CaTD_80_. Error bars represent standard error of the mean. *N* = rabbits / group, *n* = cells / group.

Having established a means to reliably evoke a cellular VP_RRC_ by pharmacological K_ATP_ activation, we sought to determine whether normal coordination of electro-mechanical dynamics – as seen in cells exposed to physiological solution (Fig. 1b, c) – protects ventricular CM from mechano-arrhythmogenesis during different phases of the AP (specifically, during late repolarisation and in diastole), and whether disruption of normal RRC increases arrhythmogenic responses to cell stretch. Acute stretch – as occurs in ischaemically weakened myocardium in the whole heart^26^ – was applied at three magnitudes with carbon fibres attached to either end of single cells under piezo-electric actuator control (Fig. 2a). Each stretch (average sarcomere stretch levels were 8 ± 1 %, 13 ± 1 %, and 17 ± 1 %) was timed from the electrical pacing stimulus to occur during the VP_RRC_ or in diastole (Fig. 2b, c). Stretch effects were assessed within one and the same cell, exposed to either physiological or pinacidil-containing solutions. Stretch resulted in a variety of changes in cellular mechanical activity (revealed by tracking sarcomere length), including stretch-induced premature contractions (sometimes followed by a second contraction out of sync with the continuing electrical stimulation; Fig. 2d and Supplementary Video 1) and other self-sustained arrhythmic responses. These sustained arrhythmic events included transient refractoriness towards electrical stimulation that resulted in single (Fig. 2e) or multiple (Fig. 2f, Supplementary Video 2) missed beat(s), or sustained activity that either resolved spontaneously (Fig. 2g) or was terminated by application of an additional stretch (Fig. 2h, Supplementary Video 3). Arrhythmia incidence was greater in pinacidil-treated cells than in control, both with stretch applied during diastole or the VP_RRC_. This finding suggests that normal electro-mechanical coordination does indeed protect CM from arrhythmogenesis during late phases of the AP, and that disturbed RRC dynamics increases overall susceptibility to mechano-arrhythmogenesis (Fig. 3a).

**Figure 2.**
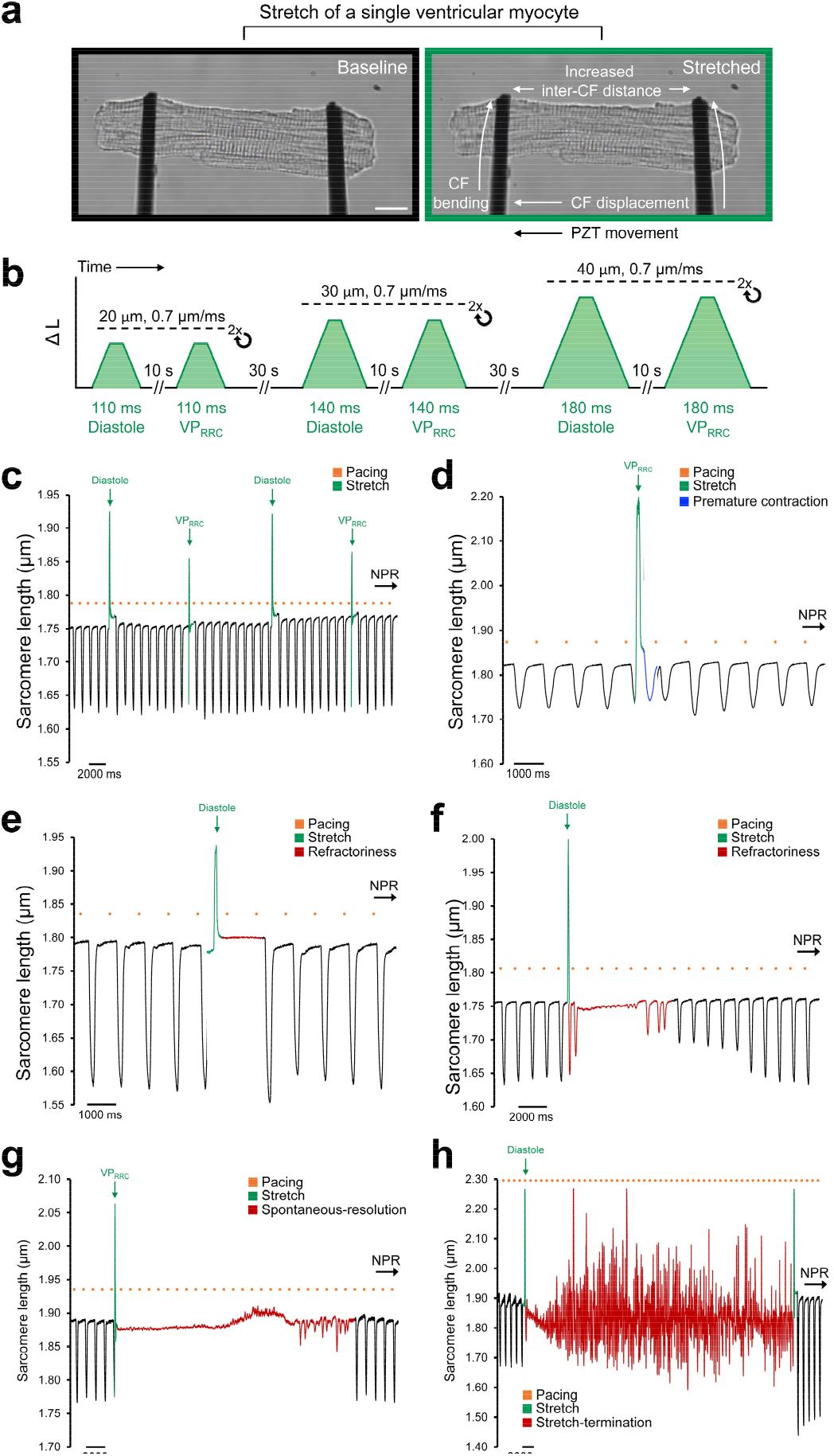
Arrhythmias elicited by acute stretch of rabbit single left ventricular cardiomyocytes. **a,** Representative brightfield image of a ventricular cardiomyocyte before (left) and during (right) axial stretch, applied using a carbon fibre-based system. Scale bar: 10 μm. **b**, Schematic of the stretch protocol. **c**, Representative measurement of sarcomere length in a cell exposed to pinacidil (50 µM, to activate ATP-sensitive potassium channels) during 1 Hz pacing (orange dots) and stretched (green segments of trace) in diastole (stretches 1 and 3) or during the vulnerable period (VP_RRC_, stretches 2 and 4), which did not result in an arrhythmia (normal paced rhythm, NPR, was maintained). **d,** Premature contraction (blue segment) upon VP_RRC_-timed stretch. **e,** Refractoriness after a diastolic stretch. **f** premature contraction after a diastolic stretch, followed by refractoriness and multiple missed beats (red segment). **g,** Self-sustained arrhythmic activity (red segment) after a VP_RRC_-timed stretch that spontaneously resolved. **h,** Self-sustained arrhythmic activity (red segment) after a diastolic stretch that was terminated by an additional stretch (second green bar).

**Figure 3.**
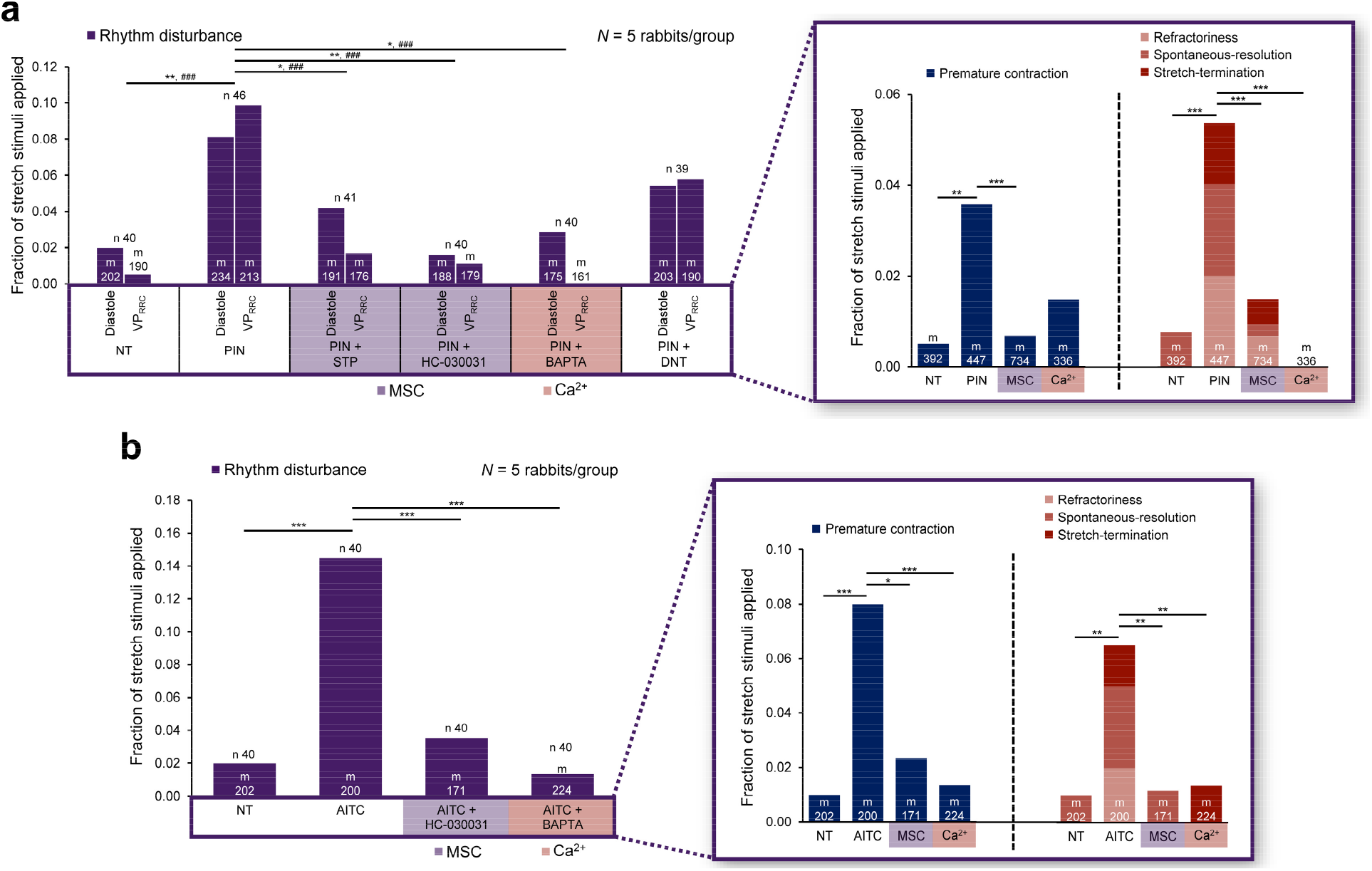
Effects of pharmacological interventions on mechano-arrhythmogenesis. **a,** Incidence of arrhythmias (purple) in rabbit isolated left ventricular cardiomyocytes with acute stretch applied during diastole or the vulnerable period (VP_RRC_) in cells exposed to one of the following treatments: Tyrode (NT); pinacidil (PIN, 50 µM, to activate ATP-sensitive potassium channels); PIN + streptomycin (STP, 50 µM, to additionally non-specifically block mechano-sensitive channels, MSC); PIN + HC-030031 (10 µM, to additionally block TRPA1 channels); PIN + BAPTA (1 µM, to additionally buffer cytosolic Ca^2+^, [Ca^2+^]_i_) or PIN + dantrolene (DNT, 1 µM, to additionally stabilise ryanodine receptors). **Inset:** of the total arrhythmia incidence (measured as the sum of diastolic and VP_RRC_ incidence), the proportion of arrhythmias that were premature contractions (blue, left) or self-sustained arrhythmic events (shades of red, right), and the effect of blocking either MSC (measured as the pooled data from stretches in the PIN + STP and PIN + HC-030031 interventions) or chelating [Ca^2+^]_i_ (measured as the pooled data from stretches in the PIN + BAPTA intervention) on their incidence in cells with activated ATP-sensitive potassium channels. Differences assessed using chi-square contingency tables and Fisher’s exact test. \**p* < 0.05, \*\**p* < 0.01, \*\*\**p* < 0.001 for diastole; #*p* < 0.05, ##*p* < 0.01, ###*p* < 0.001 for VP_RRC_. **b**, Incidence of arrhythmias with stretch during diastole in cells exposed to one of the following treatments: NT; allyl isothiocyanate (AITC, 10 µM, to activate TRPA1 channels); AITC + HC-030031; or AITC + BAPTA. **Inset:** of the total diastolic arrhythmia incidence, the proportion of those that were premature contractions (blue, left) or self-sustained arrhythmic events (shades of red, right), and the effect of blocking MSC (measured as the AITC + HC-030031 intervention) or chelating [Ca^2+^]_i_ (measured as the AITC + BAPTA intervention) on their incidence in cells with activated TRPA1 channels. Differences assessed using chi-square contingency tables and Fisher’s exact test. \**p* < 0.05, \*\**p* < 0.01, \*\*\**p* < 0.001. *N* = rabbits / group, *n* = cells / group, *m* = stretch stimuli applied.

As previous work in the whole heart had shown that the magnitude of tissue deformation is a key determinant of mechano-arrhythmogenesis,^27^ we assessed effects of stretch amplitude on arrhythmia incidence. Increasing the piezo-electric actuator displacement amplitude from 20 to 40 μm resulted in an increase in the percent sarcomere stretch (along with peak sarcomere length), as well as maximum applied stress (Extended Data Fig. 1). These increases in applied load raised the incidence of stretch-induced arrhythmias in pinacidil-treated cells, while arrhythmia incidence remained low in control cells (Extended Data Fig. 2), indicating that the increase in mechano-arrhythmogenesis that occurs with pinacidil-induced RRC disruption scales with the degree of cell stretch.

To assess whether arrhythmias could be the result of stretch-induced cellular damage, contractile function was assessed before and after stretch application of each magnitude. Both in control and in pinacidil-treated cells, diastolic sarcomere length and maximum rate and percent of sarcomere shortening were not significantly different during contractions from resting length, before and after stretch applications of any tested amplitude (Extended Data Fig. 3). These parameters were also not significantly different before and after return to steady state, following periods of self-sustained rhythm disturbances (Extended Data Fig. 4), suggesting that cellular damage may not explain mechano-arrhythmogenesis in our model.

We next sought to determine mechanisms underlying the triggering and / or sustenance of cellular mechano-arrhythmogenesis (*i.e.*, stretch-induced single excitations and self-sustained rhythm disturbances) in cells with disturbed RRC. K_ATP_ channels are normally quiescent, but if pre-activated (such as by a reduction in ATP levels during ischaemia or by pinacidil), their open probability is also increased by stretch.^28^ As K_ATP_ channels are highly selective for potassium, they have a reversal potential of about -90 mV and conduct an outward current over the entire working range of the ventricular CM membrane potential. As a result, their activation will always contribute to re-/hyperpolarisation. So, K_ATP_ channels may shorten APD, but they cannot account for stretch-induced excitation seen in this and other studies. Interestingly, while pinacidil is generally thought of as a specific activator of K_ATP_ channels, it has been shown to also increase the activity of transient receptor potential kinase ankyrin 1 (TRPA1) channels in HEK293 cells, resulting in a depolarising trans-membrane current and Ca^2+^ influx.^29^ Importantly in the context of mechano-arrhythmogenesis, TRPA1 channels: (i) are inherently mechano-sensitive^5^ and contribute to mechanically-evoked electrical responses (measured as trans-membrane currents and deduced from AP initiation) in sensory neurons,^6–8^ astrocytes,^9^ and vertebrate hair cells;^10^ (ii) have been found to be functionally expressed in murine ventricular CM^30^ (although not all studies have found this)^31^ and to drive changes in *Drosophila* heart rhythm in response to mechanical stimulation;^11^ and (iii) are important for cardiovascular function and disease progression.^13^ We hypothesised that, if TRPA1 channels are functionally expressed in rabbit left ventricular CM, then they could contribute to the increase in mechano-arrhythmogenesis seen with pinacidil treatment.

To assess the presence and function of TRPA1 channels in the rabbit left ventricle, we first measured TRPA1 protein expression in tissue from the left ventricular free wall by Western blotting, which revealed robust expression (Extended Data Fig. 5a). Next, functional expression of TRPA1 channels in left ventricular CM was evaluated by measuring ion channel activity in cell-attached patches during application of the TRPA1 channel-specific agonist, allyl isothiocyanate (AITC; 50 µM).^32^ AITC exposure caused an increase in total inward current, while there was no significant change in time-matched control cells (Extended Data Fig. 5b, c).

Having shown that TRPA1 channels are functionally expressed in rabbit ventricular CM, we explored their possible involvement in the mechano-arrhythmogenesis observed in our model. This was done by applying either a non-specific blocker of cation non-selective mechano-sensitive ion channels (streptomycin;^33^ 50 μM), or a specific TRPA1 channel blocker (HC-030031;^34^ 10 μM, which has been shown to inhibit TRPA1 current and mechanically-induced excitation in sensory neurons^6–8^) to pinacidil-treated cells. Streptomycin and HC-030031 reduced the incidence of cellular arrhythmias with pinacidil treatment, both with stretch during diastole and in the VP_RRC_ (Fig. 3a; arrhythmia incidence remained low in control cells, Extended Data Fig. 6). This finding suggests that mechano-sensitive channels are involved in mechano-arrhythmogenesis across phases of the AP when there is disruption of normal electro-mechanical coordination. More specifically, with disruption of RRC, mechano-sensitive channels appear to contribute to the increase in both single triggers and self-sustained rhythm disturbances, as their block (compared in pooled data from diastolic and VP_RRC_ stretches in cells exposed to streptomycin- or HC-030031) in pinacidil-treated cells was able to prevent the increase of both classifications of arrhythmias (Fig. 3a, inset; incidence remained low in control cells, Extended Data Fig. 6). Importantly, streptomycin and HC-030031 appeared to have no other functional effects, as there was no significant change in diastolic sarcomere length or contractile function with their application to control cells (Extended Data Fig. 7).

Next, as activation of TRPA1 channels is known to increase Ca^2+^ influx,^12, 35^ we assessed whether cytosolic Ca^2+^ contributes to the mechano-arrhythmogenesis seen in our study. This was achieved by buffering cytosolic Ca^2+^ with BAPTA-AM (1 μM, K_d_ ≍ 0.5 μM), which has previously been shown in ventricular CM to prevent stretch-induced changes in cardiac electrophysiology driven by changes in [Ca^2+^]_i_.^36^ In pinacidil treated cells, BAPTA-AM reduced arrhythmia incidence with stretch both during diastole and in the VP_RRC._ As stretch is also known to increase sarcoplasmic reticulum Ca^2+^ leak through ryanodine receptors^37^ (an effect that is enhanced in ischaemia^38^), we investigated its possible contribution to the dependence of mechano-arrhythmogenesis on cytosolic Ca^2+^ by stabilising ryanodine receptors in their closed state with dantrolene (1 μM, which has been shown to not affect Ca^2+^-induced Ca^2+^ release or contraction^39^). However, dantrolene had no significant effect on the incidence of stretch-induced arrhythmias, suggesting that Ca^2+^ influx *via* TRPA1, but not mechano-sensitive Ca^2+^ release from the sarcoplasmic reticulum, is involved in mechano-arrhythmogenesis (Fig. 3a). Yet, in contrast to mechano-sensitive channels, which contribute to both single triggers and self-sustained rhythm disturbances in pinacidil-treated cells, cytosolic Ca^2+^ appears to be involved only in the observed increase in self-sustained rhythm disturbances, as BAPTA-AM did not decrease the incidence of single excitations (compared in pooled data from diastolic and VP_RRC_ stretches in BAPTA-AM-treated cells; Fig. 3a, inset). In control cells treated with either BAPTA-AM or dantrolene, the incidence of stretch-induced arrhythmias remained low (Extended Data Fig. 6) and there was no significant change in diastolic sarcomere length or maximum rate of systolic sarcomere shortening. There was, however, a significant decrease in the percent systolic sarcomere shortening with both interventions, presumably due to a reduction in peak systolic [Ca^2+^]_i_ (by buffering with BAPTA-AM or reduced sarcoplasmic reticulum Ca^2+^ release with dantrolene; Extended Data Fig. 7).

We then sought to more specifically explore whether TRPA1 channels are indeed a source of mechano-arrhythmogenesis. Control cells were exposed to AITC (10 µM), which enhances the response of TRPA1 channels to mechanical stimulation,^8^ and then subjected to diastolic stretch (as there was no K_ATP_-induced disturbance of RRC in these cells, no VP_RRC_ stretch was applied). AITC caused an increase in both stretch-induced single excitations and self-sustained events compared to control cells (Fig. 3b). As with pinacidil exposure, application of HC-030031 to AITC-treated cells prevented this increase in mechano-arrhythmogenesis. However, unlike in pinacidil-treated cells, BAPTA-AM also prevented the increase in both types of rhythm disturbances during AITC exposure (Fig. 3b, inset).

While the above findings strongly suggest a role of TRPA1 channels in mechano-arrhythmogenesis, we did not find an increase in TRPA1 current in our patch-clamp experiments when the cell membrane within the pipette was stretched by negative pressure application. As there is a large body of evidence demonstrating that interactions between TRP channels and microtubules are important for TRP channel function,^40^ we hypothesised that microtubule-mediated mechano-transduction during axial cell stretch may be necessary for a mechanically-induced response, which would not have been engaged during local membrane stretch *via* the patch pipette. To test for the importance of microtubules in TRPA1-dependent mechano-arrhythmogenesis, cells were exposed to paclitaxel (10 µM) to increase polymerisation and stabilisation of microtubules,^41^ which has been shown to increase the probability of stretch-induced arrhythmias in isolated rabbit hearts.^42^ Immunofluorescence imaging revealed that paclitaxel increased the density of the microtubule network (Extended Data Fig. 8a-c), which was associated with an increased incidence of single excitations upon stretch in diastole (Extended Data Fig. 8d). That this increase in mechano-arrhythmogenesis involves interactions with TRPA1 channels was supported by a reduction in the incidence of stretch-induced arrhythmias in paclitaxel-treated cells during block of TRPA1 with HC-030031 (Extended Data Fig. 8d).

Our results up to this point suggest that an inward current, passing (at least in part) through mechano-sensitive TRPA1 channels, acts as a trigger for stretch-induced excitations and contributes to a substrate for self-sustained rhythm disturbances. While a transient inward depolarising current with stretch may explain the role of TRPA1 in single excitations, its contribution to the formation of an arrhythmogenic substrate for self-sustained rhythm disturbances warranted further investigation. TRPA1 channel activity has been shown to cause an increase in diastolic [Ca^2+^]_i_ in ventricular CM.^43^ This increase in diastolic [Ca^2+^]_i_ can contribute to sarcoplasmic reticulum Ca^2+^ release, creating an arrhythmogenic substrate,^44^ and can further activate TRPA1 channels.^45^ We therefore hypothesised that TRPA1-mediated effects on [Ca^2+^]_i_ may account for the observed increase in stretch-induced self-sustained rhythm disturbances. We measured [Ca^2+^]_i_ with a ratiometric fluorescence imaging technique (using the Ca^2+^-sensitive dye Fura Red, AM; K_d_ ≍ 400 nM), which showed that both pinacidil and AITC application increase [Ca^2+^]_i_ in diastole (Extended Data Fig. 9). However, despite the anti-arrhythmic effect of blocking TRPA1 channels both in AITC and in pinacidil-treated cells, HC-030031 reduced the increase in diastolic [Ca^2+^]_i_ only during AITC treatment (Extended Data Fig. 9). This finding suggests that the increase in diastolic [Ca^2+^]_i_ with AITC and pinacidil may be occurring through different mechanisms. As chelating cytosolic Ca^2+^ with BAPTA-AM was also anti-arrhythmic in both pinacidil- and AITC-treated cells (Fig. 3a, b), this suggests that increased [Ca^2+^]_i_ may be necessary for sustenance, but not sufficient for induction of mechano-arrhythmogenesis. Unfortunately, we could not measure the effects of BAPTA-AM on [Ca^2+^]_i_, as the combined buffering of Ca^2+^ by BAPTA-AM and the Ca^2+^-sensitive dye Fura Red greatly diminished cell contraction.

Explicit evidence for a direct role of Ca^2+^ in the self-sustained rhythm disturbances came from dual voltage-Ca^2+^ fluorescence imaging, which revealed an increase in [Ca^2+^]_i_, along with Ca^2+^ oscillations that precede changes in voltage, indicating that oscillations in [Ca^2+^]_i_ can drive sustained aberrant electrical behaviour (Fig. 4a, Supplementary Video 4). It is worth noting here that changes in repolarisation can also destabilise the electrical activity of ventricular CM. Thus, in pinacidil-treated cells, a stretch-induced increase in K_ATP_ channel current may have directly contributed to the ability to sustain arrhythmic activity by shortening APD, thereby increasing the diastolic interval and the duration of the excitable period between paced beats.^28^

**Figure 4.**
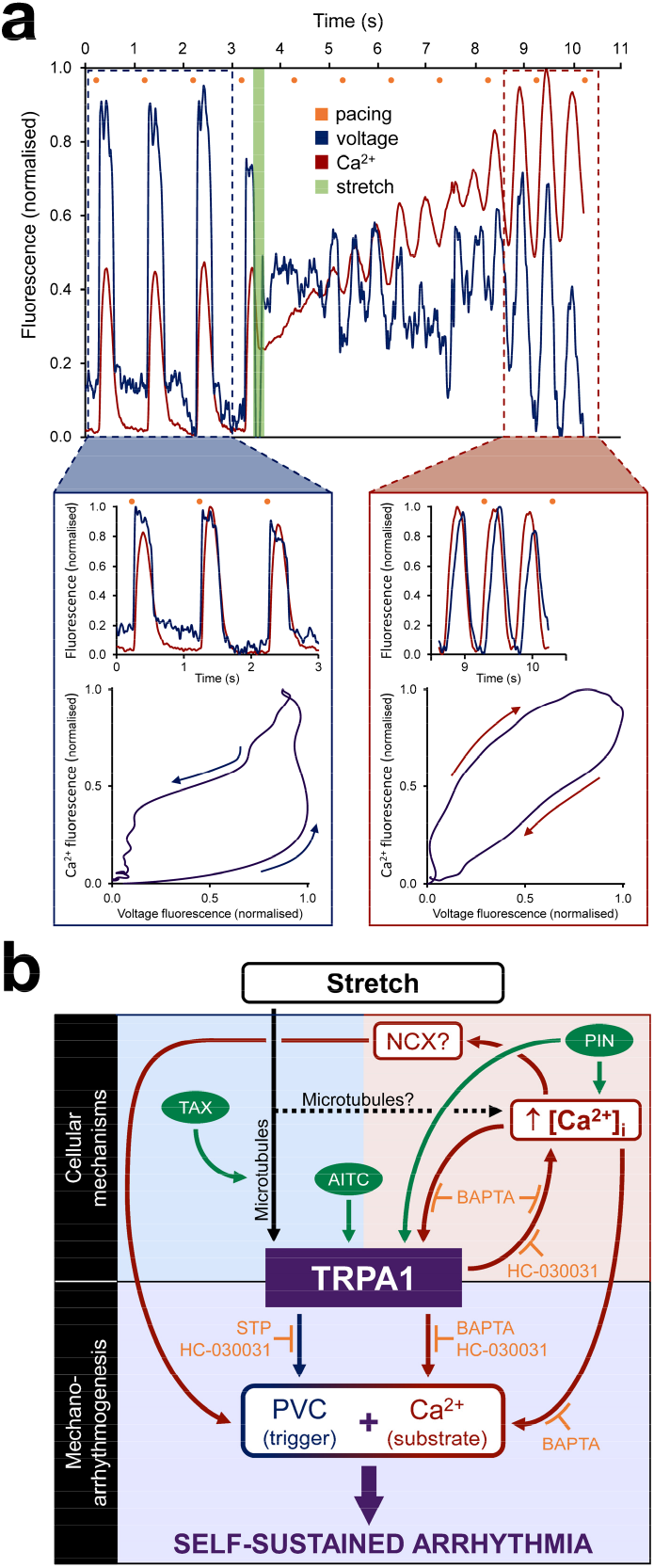
Role of TRPA1 channels and cytosolic calcium ([Ca^2+^]_i_) in ventricular mechano-arrhythmogenesis. **a,** Example trace of voltage (blue) and [Ca^2+^]_i_ (red), simultaneously recorded by fluorescence imaging in a rabbit isolated left ventricular cardiomyocyte exposed to pinacidil (PIN, 50 µM, to activate K_ATP_ channels). Rapid stretch in the vulnerable period (VP_RRC_, green) resulted in sustained arrhythmic activity, initiated by sustained depolarisation and an associated increase in [Ca^2+^]_i_ above normal systolic levels, prompting spontaneous oscillations in [Ca^2+^]_i_ (which subsequently resolved, followed by normal paced beats; not shown). **Blue inset**: Scaled voltage and Ca^2+^ fluorescence curves (top) and phase plot of the first beat (bottom) showing that, prior to stretch, changes in voltage preceded Ca^2+^. **Red inset**: Scaled voltage and Ca^2+^ fluorescence curves (top) and phase plot of the first beat (bottom) showing [Ca^2+^]_i_ oscillations preceding changes in voltage during sustained arrhythmia after stretch, suggesting aberrant Ca^2+^ handling as a driving mechanism. **b,** Schematic representation of mechanisms underlying TRPA1 channel driven mechano-arrhythmogenesis. Effects of stretch (black) on cell electrophysiology (blue), Ca^2+^ (red), or a combination of stretch and Ca^2+^ (purple). TRPA1 agonists (green) and targets of pharmacological antagonists (orange) are indicated. AITC, Allyl isothiocyanate; NCX, Na^+^-Ca^2+^ exchanger; PIN, pinacidil; STP, streptomycin; TAX, paclitaxel.

Finally, with an increase in [Ca^2+^]_i_, there is the possibility that a change in cellular mechanics may also contribute to mechano-arrhythmogenesis by enhancing mechano-transduction or altering the characteristics of stretch. However, we observed no significant effect of pinacidil on diastolic or systolic cell stiffness, measured (respectively) as the slopes of end-diastolic and end-systolic stress-sarcomere length relationship in mechanically loaded (*via* carbon fibres) contracting cells (Extended Data Fig. 10). Furthermore, we found no significant effect of pinacidil on percent sarcomere stretch, stretched sarcomere length, or maximum applied stress during stretch (Extended Data Fig. 1). This suggests that changes in cellular mechanics may not be a major contributor to observed effects (note: there was a significant shift in the stiffness curves to the right in pinacidil-treated cells, which may relate to reduced affinity of myofilaments for Ca^2+^ through TRPA1-induced phosphorylation of troponin I;^43^ Extended Data Fig. 10). In a subset of pinacidil-treated cells for which two consecutive stretches of the same magnitude (occurring either during diastole or the VP_RRC_) first did, and then did not, result in an arrhythmia, the characteristic effects of each stretch were not significantly different (Extended Data Fig. 11), suggesting that passive cell mechanical properties were not affected by acute stretch application.

Taken together, our results suggest that TRPA1 channels contribute both to acute triggering and to Ca^2+^-mediated sustenance of ventricular mechano-arrhythmogenesis. Figure 4b summarises our findings and presents a working model of TRPA1’s role in triggering stretch-induced excitation (through a depolarising trans-sarcolemmal current) and in creating a substrate for sustained arrhythmic activity (through increased diastolic [Ca^2+^]_i_).

TRPA1 channels are mechano-sensitive and thus allow for a trans-sarcolemmal influx of cations upon stretch (specifically Ca^2+^ when TRPA1 channels are pre-activated),^12, 35^ which depolarises membrane potential. An increase in [Ca^2+^]_i_ may result in further membrane depolarisation *via* electrogenic forward-mode sodium-Ca^2+^ exchanger activity.^44^ If the depolarisation is sufficiently large, it will cause excitation and premature contraction of the cell, with arrhythmic activity sustained if [Ca^2+^]_i_ remains sufficiently elevated for a period of time. Importantly, TRPA1 channel activity is directly modulated by [Ca^2+^]_i_. Specifically, an increase in [Ca^2+^]_i_ potentiates TRPA1 channel activity, but at a certain level of [Ca^2+^]_i_ TRPA1 channels are progressively inactivated, thus limiting the degree of Ca^2+^ overload and, perhaps, preventing lethal cell damage.^46^ In our experiments, exposure to AITC caused direct activation of TRPA1 channels and subsequent diastolic [Ca^2+^]_i_ loading, while pinacidil either directly activated TRPA1 channels, or did so indirectly *via* an increase in [Ca^2+^]_i_.

In conclusion, our results suggest that TRPA1 activation leads to enhanced microtubule-dependent, stretch-induced depolarising currents and increased [Ca^2+^]_i_. As such, block of TRPA1 channels reduces both the triggering and self-sustenance of mechano-arrhythmogenesis, while buffering diastolic [Ca^2+^]_i_ only reduces arrhythmia sustenance. These findings have potentially important implications for anti-arrhythmic strategies in cardiac pathologies associated with augmented TRPA1 channel expression or activity.^13^ This is especially true for cases in which additional factors activate TRPA1 channels (*e.g.*, oxidative stress),^47^ as the response of TRPA1 to mechanical stimulation is enhanced by an increase in their baseline activity.^8^ For instance, acute myocardial ischaemia is associated with TRPA1-mediated myocardial damage^14^ (which can be reduced by TRPA1 inhibition^15^), along with lethal ventricular arrhythmias that are thought to involve contributions of altered tissue mechanics, intracellular Ca^2+^ handling, and oxidative stress.^26^ Thus, targeting TRPA1 channels in that setting may help in preventing multiple detrimental outcomes. This could also be true in chronic pathologies in which lethal arrhythmias occur, and which are associated with changes in cardiac mechanics, intracellular Ca^2+^ dynamics, oxidative stress, and other TRPA1 modulating factors,^12^ such as ventricular pressure overload. In keeping with this suggestion, TRPA1 inhibition in chronic hypertension has been shown to reduce hypertrophy and fibrosis.^16^ Overall, these observations, combined with our current findings, suggest TRPA1 channels may represent a novel, underappreciated anti-arrhythmic target with exciting therapeutic potential.^17^

## METHODS

### Animal model

Experiments involved the use of female New Zealand White rabbits (2.1 ± 0.2 kg, Charles River) – the most relevant small animal model for cardiac arrhythmia research^48^ – and were conducted in accordance with the ethical guidelines of the Canadian Council on Animal Care as well as the German legislation for animal welfare, with protocols approved by the Dalhousie University Committee for Laboratory Animals, or by the local Institutional Animal Care and Use Committee at the University of Freiburg (Regierungspräsidium Freiburg, X-16/10R). Details have been reported following the Minimum Information about a Cardiac Electrophysiology Experiment (MICEE) reporting standard.^49^

### Cell isolation

Animals were euthanised by ear vein injection of pentobarbital (140 mg/kg) and heparin (1,500 units/kg, Sigma-Aldrich), followed by rapid excision of the heart, aortic cannulation, and Langendorff perfusion (20 mL/min) with 37 °C modified Tyrode (NT) solution (containing, in mM: 120 NaCl, 4.7 KCl, 1 MgCl_2_, 1.8 CaCl_2_, 10 glucose, 10 HEPES [Sigma-Aldrich], with pH adjusted to 7.40 ± 0.05 with NaOH and an osmolality of 300 ± 5 mOsm/L) bubbled with 100 % oxygen. After a rest period of 15 min, the perfusate was switched to calcium (Ca^2+^)-free cardioplegic solution for 5 min (containing, in mM: 117 NaCl, 10 KCl, 1 MgCl_2_, 10 creatine, 20 taurine, 5 adenosine, 2 L-carnitine, 10 glucose, 10 HEPES [Sigma-Aldrich], with pH adjusted to 7.40 ± 0.05 with NaOH and an osmolality of 300 ± 5 mOsm/L), with an addition of 0.018 mM EGTA (Sigma-Aldrich). The perfusate was then switched to digestion solution, comprised of Ca^2+^-free solution with the addition of 200 U/mL Collagenase II (Worthington Biochemical Corporation), 0.06 mg/mL Protease XIV (from *Streptomyces griseus*, Sigma Aldrich), and 100 µM CaCl_2_. After 5 min, the perfusion rate was reduced to 15 mL/min until the heart became flaccid and jelly-like (∼10-12 min). The left ventricular free wall was removed and placed into 50 mL of stop solution, comprised of the Ca^2+^-free cardioplegic solution described above without EGTA, but with an addition of 0.5 % bovine serum albumin (Sigma Aldrich) and 100 µM CaCl_2_. The tissue was agitated, the solution filtered through a 300 µm pore nylon mesh, the tissue returned to fresh stop solution and re-agitated, and the solution filtered in duplicate. The cell-containing filtered solution was divided into 2 mL microcentrifuge tubes and maintained at room temperature.

### Carbon fibre method

Cells were stretched using the carbon fibre method, adapted from previous work.^19^ Briefly, single carbon fibres (12-14 µm in diameter,) were mounted with cyanoacrylate adhesive into a pair of borosilicate glass capillaries, pulled from glass tubes (1.12 mm inner / 2 mm outer diameter, World Precision Instruments) and bent at 30 **°** to allow near-parallel alignment of the fibres with the bottom of the experimental chamber. Carbon fibres were trimmed to a length of either 1.2 mm or 0.6 mm from the end of the glass capillary and used in combination so that one fibre was relatively compliant and the other relatively stiff. Each carbon fibre-containing capillary was mounted in a microelectrode holder (MEH820, World Precision Instruments), coupled to a three-axis water hydraulic micromanipulator (MHW-103, Narishige) on top of a linear piezo-electric translator (PZT, P-621.1CD, Physik Instrumente). The carbon fibre position was controlled by a piezo amplifier / servo controller (E-665.CR, Physik Instrumente), driven by a voltage signal generated from a data acquisition (DAQ) device (USB-6361; National Instruments) and custom-written routines in LabVIEW (National Instruments). Prior to use on CM, carbon fibre stiffness was calibrated by pressing a fibre against a force transducer system (406A, Aurora Scientific), measuring the force for given displacements of the PZT, and fitting the data by linear regression to the formula: stiffness = force / carbon fibre bending (where bending = PZT displacement). This allowed measurement of passive and active forces of CM by tracking carbon fibre bending (monitoring fibre tips and prescribing PZT position; described below). Force was calculated as: force = stiffness × (change in distance between carbon fibre tips - change in distance between PZT positions). Stress was assessed by dividing force by the stretched cross-sectional area (CSA) of the CM, determined by assuming that the cross section is an ellipse with CSA = π × (cell width / 2) × (cell thickness / 2), where cell thickness is assumed to be = width / 3.^50^

### Single cell stretch

Before use, half of the supernatant from the cell-containing microcentrifuge tube was removed and replaced by room temperature NT for 10 min (so that cells were exposed to 50 % of normal extracellular Ca^2+^ concentration), after which all of the supernatant was removed and replaced by NT. A drop of cell-containing solution was added to an imaging chamber (RC-27NE2, Warner Instruments) containing 1 mL of NT maintained at 35 °C by a temperature controller (TC-344C, Warner Instruments) and mounted on an inverted fluorescence microscope (IX-73, Olympus) with a 40× objective (UPLFLN40X, Olympus). The surface of the coverslip on the bottom of the chamber was coated with poly-2-hydroxyethyl methacrylate (poly-HEMA, Sigma-Aldrich) to prevent cell adhesion. Once the cells had settled, 1 Hz biphasic electrical field stimulation (100 mA, 2 ms square pulse duration; SIU-102, Warner Instruments) was commenced and homogeneously contracting, rod-shaped CM with clear striations and well-defined membranes lacking signs of blebbing were selected. Electrical stimulation was stopped, and one long and one short carbon fibre were positioned above either end of a CM and gently lowered onto the cell surface using the hydraulic micromanipulators. Adhesion of the CM to carbon fibres was confirmed by raising the cell off the coverslip. Once cell attachment was established, electrical stimulation was recommenced, and cells contracted against the mechanical afterload imposed by carbon fibres for ∼1-2 min, which improved cell adhesion. Cells were then superfused at 2.1 mL/min through an inline heater (SF-28, Warner Instruments) with either NT or pinacidil-containing solution (50 µM, to activate ATP-sensitive potassium, K_ATP_, channels) for 5 min. To approximate stretch of ischaemically weakened myocardium during acute regional ischaemia,^51–55^ transient stretch (20 µm PZT displacement, applied and removed at a rate of 0.7 μm/ms) was applied (Fig. 2a). Stretch was timed from the electrical stimulus, to occur at two times during the AP, with 10 s between each stretch: first during mid-diastole (600 ms delay after an electrical stimulus), and then during the vulnerable period of the repolarisation-relaxation coupling interval (VP_RRC_; 150 ms delay, so that a majority of stretches – whose duration increased with greater PZT displacement – would occur within the VP_RRC_, based on the timing measured by simultaneous voltage-Ca^2+^ imaging; described below). These two differentially timed stretches were then duplicated, such that there was a total of four stretches at each magnitude of PZT displacement. This protocol was repeated with increasing magnitudes of PZT displacement (30 and 40 µm) to generate a range of stretch-induced changes in sarcomere length within a cell (30 s between repetitions, for a total of 12 stretches; Fig. 2b).

### Assessment of stretch effects

Throughout the protocol, sarcomere length, PZT location, and carbon fibre tip positions were monitored and recorded at 240 Hz (Myocyte Contractility Recording System, IonOptix). From this, CM contractile function (diastolic sarcomere length, and maximum rate and percent sarcomere shortening), as well as characteristics of cell stretch (percent sarcomere stretch, peak sarcomere length, and maximum applied stress) were obtained. Arrhythmic activity was assessed from sarcomere length measurements, which revealed stretch-induced premature contractions (defined as 1 or 2 unstimulated contractions after stretch), or other self-sustained arrhythmic activity (including refractoriness, resulting in single or multiple missed beats, or sustained activity that either spontaneously resolved, or was terminated by an additional stretch). Examples of each type of stretch-induced arrhythmia are provided in Fig. 2. Any stretch that resulted in carbon fibre detachment, or a sustained arrhythmia that did not spontaneously resolve or could not be terminated by a maximum of two additional stretch applications was excluded (< 1 % of all cells).

### Pharmacological interventions

Pharmacological agents dissolved as stock solutions in distilled water, dimethyl sulfoxide (DMSO), or ethanol, as appropriate, were added to the perfusate and continuously perfused for 5 min before any recordings. Agents included: BAPTA-AM (1 µM from a stock concentration of 10 mM in DMSO, to buffer cytosolic Ca^2+^, with the concentration determined in preliminary experiments by titrating to a value that caused a ∼ 10 % decrease in percent sarcomere shortening during contraction, Abcam), dantrolene (1 µM from a stock concentration of 5 mM in DMSO, to stabilise ryanodine receptors, Abcam), streptomycin (50 µM from a stock concentration of 15 mM in distilled water, to non-specifically block mechano-sensitive channels, Sigma-Aldrich), HC-030031 (10 µM from a stock concentration of 50 mM in DMSO, to block TRPA1 channels, Abcam), allyl isothiocyanate (AITC, 10 µM from a stock concentration of 1 M in DMSO, to activate TRPA1 channels, Sigma-Aldrich), and paclitaxel (10 µM from a stock concentration of 10 mM in DMSO, to increase the polymerisation and stabilisation of microtubules, Abcam). For BAPTA-AM, HC-030031, and paclitaxel, cells were incubated for 20, 30, or 90 min respectively, after introduction to NT but prior to superfusion.

### Dual parametric voltage-Ca^2+^ fluorescence imaging

The Ca^2+^-sensitive dye Fluo-5F, AM (5 µM from a stock concentration of 1 mM in DMSO, ThermoFisher Scientific) and Pluronic F-127 (0.02 % from a stock concentration of 20 % in DMSO, Biotium) were added to the cell-containing microcentrifuge tube during cellular suspension in the 50 % Ca^2+^ solution, and cells were incubated in the dark for 20 min. The supernatant was then replaced with fresh NT, the voltage-sensitive dye di-4-ANBDQPQ (20 μM from a stock concentration of 28.8 mM in ethanol, University of Connecticut Health Centre) was added to the tube, and the cells were incubated in the dark for 14 min. The supernatant was again replaced with fresh NT, probenecid (1 mM from a stock concentration of 250 mM in distilled water, Sigma-Aldrich) was added to the tube, and the cells were from there on maintained in the dark at room temperature. For imaging, the solution containing dye-loaded cells was gently agitated with a transfer pipette and a drop of solution was added to 1 mL of NT in the imaging chamber. Carbon fibres were attached to a cell as described above (carbon fibres reduced motion artefact associated with cell translocation in the vertical plane during contractions, allowing measurements to be made without an excitation-contraction uncoupler). Fluorescence was excited by light from a mercury lamp (U-HGLGPS, Olympus), passed through a 466 / 40 nm bandpass filter (FF01-466/40, Semrock) and reflected onto the sample by a 495 nm dichroic mirror (FF495-Di03, Semrock). For simultaneous measurement of trans-membrane voltage and cytosolic ‘free’ Ca^2+^ concentration ([Ca^2+^]_i_), each fluorescent signal was projected onto one-half of a 128 × 128-pixel, 16-bit electron-multiplying charge-coupled device (EMCCD) camera sensor (iXon3, Andor Technology) using an emission image splitter (Optosplit II; Cairn Research), and recorded at 500 fps with 2 ms exposure time and maximum electron-multiplying gain. The two signals were split with a 685 nm dichroic mirror (FF685-Di02, Semrock) and Fluo-5F emission was collected with a 525 / 50 nm bandpass filter (FF03-525 / 50, Semrock) and di-4-ANBDQPQ emission with a 700 nm long-pass filter (HQ700lp; Chroma Technology). A schematic of the imaging setup is provided in Fig. 1a.

Analysis of voltage and [Ca^2+^]_i_ signals was performed using custom routines in Matlab (R2018a, MathWorks). Fluorescence for each signal was averaged over the entire cell, a temporal filter (50 Hz low-pass Butterworth) was applied, and effects of bleaching were corrected by fitting the resulting signal with a second-order polynomial and subtracting the result. From the corrected signals, time to 50 % or 80 % recovery of the action potential (action potential duration, APD_50_ or APD_80_) or the [Ca^2+^]_i_ transient ([Ca^2+^]_i_ transient duration, CaTD_50_ or CaTD_80_) were averaged over 3 consecutive cardiac cycles. The VP_RRC_ was calculated as the difference between CaTD_80_ and APD_50_, plus the difference between the timing of the action potential and [Ca^2+^]_i_ transient upstrokes, measured at the point of maximum upstroke velocity for each signal (excitation-contraction coupling time, ECC, see Fig. 1b, d): VP_RRC_ = (CaTD_80_ - APD_50_) + ECC.

### Ratiometric [Ca^2+^]_i_ fluorescence imaging

Imaging of [Ca^2+^]_i_ levels was performed using the ratiometric Ca^2+^-sensitive dye Fura Red, AM (K_d_ ≍ 400 nM, AAT Bioquest). Cells were incubated in the dark for 20 min with 5 µM of the dye (from a stock concentration of 1 mM in DMSO), 0.02 % Pluronic F-127, and 1 mM probenecid. Fluorescence was excited by alternating light pulses from two white light-emitting diodes (CFT-90-W; Luminus Devices), one with a 420 / 10 nm bandpass filter (FF01-420 / 10, Semrock) and the other with a 531 / 22 nm bandpass filter (FF02-531/22, Semrock), which were combined into the microscope excitation light path with a 455 nm dichroic mirror (AT455dc, Chroma Technology) and reflected onto the sample by a 562 nm dichroic mirror (T562lpxr, Chroma Technology). Fluorescence was measured through a 632 / 60 nm bandpass filter (AT635 / 60m, Chroma Technology) with the EMCCD camera at 500 fps with 2 ms exposure time and maximum electron-multiplying gain. Light pulses and camera frame acquisition were synchronised with a custom control box (provided by Dr. Ilija Uzelac, Georgia Institute of Technology) so that alternating frames corresponded to the signal generated by each of the two excitation wavelengths.^56^

Analysis of [Ca^2+^]_i_ was performed using custom routines in Matlab. Fluorescence was averaged over the entire cell and a temporal filter (50 Hz low-pass Butterworth) was applied. The two Ca^2+^ signal emissions were separated, and the ratio of the signals was calculated. Any remaining bleaching was corrected by fitting the resulting signal with a second-order polynomial and subtracting the result. From the corrected signals, the minimum value for each cardiac cycle (representing a relative measurement of diastolic [Ca^2+^]_i_) was established and averaged over 3 consecutive cardiac cycles.

### Cell immunofluorescence

Coverslips (22 mm round; VWR) were coated with laminin (100 µg/mL from Engelbreth-Holm-Swarm murine sarcoma basement membrane diluted in PBS, Sigma-Aldrich) and stored in a 12-well plate (VWR). Isolated CM in NT were added to each well and maintained at room temperature for 3 hours to seed. In half of the wells, paclitaxel (10 µM) was added for the final 90 min. Cells were then fixed by removing the NT and submerging the cover slips in ice cold methanol and incubating for 7 min at −20 °C. Methanol was removed and cover slips were rinsed 5 times with phosphate buffered saline (PBS, Sigma-Aldrich) to remove excess methanol. Cover slips were stored in PBS until staining.

For staining, coverslips were bathed in blocking solution (5 % BSA in PBS, Sigma-Aldrich) at room temperature for 1 hour. Coverslips were then transferred to a custom-built humidity chamber (a glass Petri dish lined with a PBS-soaked piece of tissue placed on upside down on top of a piece of Parafilm). Diluted primary antibody (100 µL of 1:200 rat anti-rabbit a-tubulin, Clone YL1 / 2, Invitrogen) was pipetted onto the Parafilm and the coverslips were placed cell-side down on top of the solution and incubated at 4 °C for 1 hour. Coverslips were subsequently washed in quadruplicate (5 min each) with PBS and diluted secondary antibody (1:500 goat anti-rat IgG Alexa Fluor 488 conjugate, Invitrogen) was applied as above. Following secondary antibody staining, coverslips were mounted cell-side down to glass slides with mounting medium (ProLong Glass Antifade Mountant with NucBlue Stain, ThermoFisher).

Imaging of fixed and stained CM was performed on an inverted confocal microscope system (TCS SP8, Leica) with a 40x oil immersion, 1.3 NA objective (HC Plan APOCHROMAT CS2, Leica) and Lightning deconvolution software to obtain super-resolution images. Samples were illuminated with a 488 nm solid state laser at 1.2 % intensity and the photomultiplier tube (PMT) detectors were set to collect between 503-577 nm. A z-stack of 3 images, spaced at 1.5 μm, was obtained. Maximum projections were generated using Leica LAS X 3D viewer software. Images were then analysed in a blinded fashion, using custom software in Fiji (kindly shared by Drs Matthew Caporizzo and Benjamin Prosser, Department of Physiology, University of Pennsylvania Perelman School of Medicine, Philadelphia, USA) to determine microtubule density, calculated as the fraction of cell area.^57^ Briefly, cells were manually traced and a threshold for microtubule positive pixels was determined from a background region within the cell with no visible microtubule fluorescence. This was used to generate a binary image of the cell to calculate the microtubule positive fraction of the total cell area.

### Tissue Western blotting

Boiled samples of left ventricular free wall (20 μg) were separated (4 % stacking gel) and resolved *via* 7.5 % sodium dodecyl sulfate-polyacrylamide gel electrophoresis (Mini-PROTEAN SDS-PAGE, BioRad). Samples were loaded alongside 10 µL protein ladder (Precision Plus Protein Standards Kaleidoscope ladder, BioRad). Self-cast gels (Mini-PROTEAN, BioRad) were run on ice at 90 V for 30 min and then at 120 V until the dye front migrated to the opposite edge of the gel on ice in 1X Tris / Glycine / SDS electrophoresis buffer (BioRad). Samples were wet-transferred to nitrocellulose (0.2 µm, BioRad) at 100 V for 1.5 hours buried in ice. Membranes were briefly rinsed in double-distilled water and equal protein transfer was confirmed by incubating the membranes in stain (Pierce Reversible Memcode Stain, Thermo Scientific) for 5 min. The stained blot was labelled and imaged (ChemiDoc MP Imaging System, BioRad) before removing the stain (Pierce Stain Eraser, Thermo Scientific). Membranes were then blocked in 5 % skim-milk in 1X Tris-Buffered-Saline-Tween 20 (TBS-T, Sigma-Aldrich) for 60 min. Membranes were subsequently incubated overnight at 4 °C with primary antibodies (1:2,000 monoclonal mouse anti-TRPA1 antibody, Sigma-Aldrich, and 1:3,000 MF20 mouse anti-myosin heavy chain antibody, Hybridoma Bank), diluted in 1 % skim-milk in TBS-T with sodium azide. Next, blots were incubated with secondary antibody (1:4000 horseradish peroxidase HRP-conjugated anti-mouse produced in goat, Jackson Labs) for 1 hour in 5 % skim-milk at room temperature. Immunoreactivity was then measured (Clarity Western Enhanced Chemiluminescence Substrate, with a ChemiDoc MP Imaging System, BioRad). Membranes were stripped by incubation in 25 mL 0.5 M Tris-HCl / SDS buffer supplemented with 125 μL 2-mercaptoethanol (Sigma-Aldrich) for 1 hour and re-probed.

### Patch-clamp current recordings

Cells for patch clamp experiments were isolated from ventricular slices according to a previously published protocol.^58^ Briefly, animals were anaesthetised by intramuscular injection of esketamine hydrochloride (0.5 mL/kg) and 2 % xylazine hydrochloride (0.2 mL/kg). Under sedation, an anaesthetic mixture of sodium-heparin (1,000 units/mL) and esketamine hydrochloride (25 mg/mL) were injected intravenously. Euthanasia was induced by intravenous injection of sodium-thiopental (25 mg/mL) until apnoea. The heart was then swiftly excised, cannulated, and Langendorff perfused (20 mL/min) for 2 min with 37 °C modified NT solution (containing, in mM: 137 NaCl, 4 KCl, 10 MgCl_2_, 1.8 CaCl_2_, 10 glucose, 10 HEPES, 10 creatine, 20 taurine, 5 adenosine, 2 L-carnitine; with pH adjusted to 7.30 ± 0.05 with NaOH) bubbled with 100 % oxygen, followed by 2 min with 37 °C cutting solution (containing, in mM: 138 NaCl, 5.4 KCl, 0.33 NaH_2_PO_4_; 2.0 MgCl_2_, 0.5 CaCl_2_, 10 HEPES, 30 2,3-butanedione 2-monoxime (BDM); with pH adjusted to 7.30 ± 0.05 with NaOH) bubbled with 100 % oxygen. The left ventricular free wall was cut into approximately 0.8 mm x 0.8 mm wide transmural tissue chunks, embedded in 4 % low melting point agarose dissolved in cutting solution. Left ventricular blocks were cut into 300 μm thick live tissue slices, using a vibrating microtome (7000 smz-2, Campden Instruments Ltd) with stainless-steel blades (7550-1-SS, Campden Instruments Ltd) at 4 °C, with 60 Hz horizontal blade displacements of 1.5 mm, at a blade progression velocity of 0.05 mm/s (Z-deflection values were below 1 µm). Slices were stored at 4 °C in cutting solution until digestion.

Slices were digested at 37 °C in Petri dishes (Ø 35 mm) placed on a heated orbital shaker (50-65 rpm). Slices were washed three times (1 min for each wash) with the solution used for cell isolation, but without digestive enzymes (containing, in mM: 100 NaCl, 15 KCl, 2.5 KH_2_PO_4_, 2 MgCl_2_, 10 glucose, 10 HEPES, 20 taurine, 20 L-glutamic acid monopotassium salt, 30 BDM; with 2 mg/mL bovine serum albumin; pH adjusted to 7.30 ± 0.05 with NaOH). Slices were then digested for 12 min by including proteinase type XXIV (0.5 mg/mL; Sigma-Aldrich) to the cell isolation solution mentioned above, followed by three more washes. Digestion was continued by adding 200 μL of Liberase^TM^ TL Research Grade (0.25 mg/mL; Hoffmann-La Roche) per 2 mL of cell isolation solution, supplemented with 5 μM CaCl_2_. When slices started to show a corrugated appearance (∼ 25 - 50 min), they were washed three times with the solution used for cell isolation, without enzymes but containing 10 mg/mL bovine serum albumin, and mechanically dissociated using forceps and gentle pipetting with Pasteur pipettes. The Ca^2+^ concentration was increased in a stepwise fashion every 5 min (to 5 μM, 12 μM, 20 μM, 42 μM, 84 μM, 162 μM, 266 μM, 466 μM, 827 μM, and 1 mM), during which time BDM was gradually decreased to a final concentration of 13 mM. Cells were filtered through a 1-mm-pore nylon mesh and allowed to settle in a conical centrifuge tube for 15 - 20 min. The supernatant was removed and replaced by modified NT. Quality of the preparation was assessed by the percentage of rod-shaped cells, the response to electrical pacing, and the resting sarcomere length. Cells with irregular / unclear sarcolemma, blebs, or unclear striations were rejected.

Sarcolemmal ion channel activity was recorded at room temperature (21 ± 2 °C) using the patch-clamp technique with a holding potential of +40 mV. Recordings were obtained in cell-attached configuration using a patch-clamp amplifier (Axopatch 200B, Axon Instruments) and a digitiser interface (Axon Digidata 1440A, Axon Instruments). Fire-polished soda-lime glass capillaries (1.15 ± 0.05 mm inner / 1.55 ± 0.05 mm outer diameter; VITREX Medical A/S) were pulled using a two-stage pipette-puller (PC-10, Narishige) to create the micropipettes required for patch-clamping. Average pipette resistance was 1.36 ± 0.10 MΩ. Currents were acquired at 20 kHz sampling rate (interval 50 µs), and low-pass filtered at 1 kHz. Currents were analysed with pCLAMP 10.6 software (Axon Instruments). The bath solution contained (in mM): 155 KCl, 3 MgCl_2_, 5 EGTA, 10 HEPES (Sigma-Aldrich), with pH adjusted to 7.2 ± 0.05 using KOH and an osmolality of 310 ± 5 mOsm/L. The solution was stored at room temperature. Pipette solution for cell-attached recordings contained (in mM): 150 NaCl, 5 KCl, 10 HEPES, 2 CaCl_2_ with pH adjusted to 7.4 ± 0.05 using NaOH and an osmolality of 300 ± 5 mOsm/L. The pipette solution also contained 10 mM tetraethylammonium chloride (TEA), 5 mM 4-aminopyridine, and 10 mM glibenclamide to inhibit potentially contaminating potassium channels. These ionic conditions were used previously to assess cation non-selective channels.^59^ AITC was added to the bath solution before use and perfused *via* a local perfusion system. Flow rate was adapted by adjusting the height of reservoirs for gravity-fed flow to 1 mL/min. Recordings were analysed in Clampfit 10.6. Average current was calculated over at least 10 s for each condition.

### Statistics

Statistics were performed using Prism 9 (GraphPad). Differences in arrhythmia incidence were assessed using chi-square contingency tables and Fisher’s exact test. Data was first tested for normality, then differences between group means were assessed by two-tailed, paired or unpaired Student’s t-test (for normally distributed data), Wilcoxon matched-pairs (paired) or Mann-Whitney (unpaired) test (for data that was not normally distributed), one-way ANOVA with Tukey *post-hoc* tests (for normally distributed data), or Kruskal-Wallis with Dunn’s multiple comparisons test (for data that was not normally distributed), where appropriate. The relevant test is indicated in the figure captions. A *p*-value of < 0.05 was considered indicative of a significant difference between observed parameters. Figures indicate the number of replicates used in each experiment (*N* = rabbit hearts / group, *n* = cells / group, *m* = stretch stimuli applied / analysed).

### Data availability

The datasets obtained and analysed in the current study are available from the corresponding author upon reasonable request.

### Code availability

All custom computer source code used in this study is available from the corresponding author upon reasonable request.

## STATEMENTS

### Acknowledgements

We thank Gentaro Iribe (Asahikawa Medical University, Asahikawa, Japan) and Keiko Kaihara (Okayama University, Okayama, Japan) for technical assistance with cell stretch. Carbon fibres were a gift from Jean-Yves LeGuennec. This work was supported by the Canadian Institutes of Health Research (MOP 342562 to T.A.Q.); by the Natural Sciences and Engineering Research Council of Canada (RGPIN-2016-04879 to T.A.Q.); by the Dalhousie Medical Research Foundation (T.A.Q.); by the Canadian Foundation for Innovation (32962 to T.A.Q.); and by the Heart and Stroke Foundation of Canada through a National New Investigator award (T.A.Q.). B.A.C., J.G., R.P. and P.K. are members of the German Research Foundation Collaborative Research Centre SFB1425 (DFG #422681845).

### Author contributions

B.A.C. and T.A.Q. designed the study, interpreted the data, and wrote the manuscript; B.A.C. performed and analysed the cell stretch, stiffness, and fluorescence imaging experiments, and isolated cells for the immunofluorescence experiments; M.R.S. contributed to experimental design and setup; J.G. isolated cells for the patch-clamp experiments; R.P. performed and analysed the patch-clamp experiments; M.S.C. performed and analysed the Western blotting experiments; and J.J.B. and E.A.D. performed and analysed the immunofluorescence experiments. P.K. contributed to the intellectual content of the research and revised the manuscript. All authors read and approved the manuscript.

### Author information

The authors declare no competing interests. Correspondence and request for materials should be addressed to T.A.Q.

## EXTENDED DATA

**Extended Data Figure 1.**
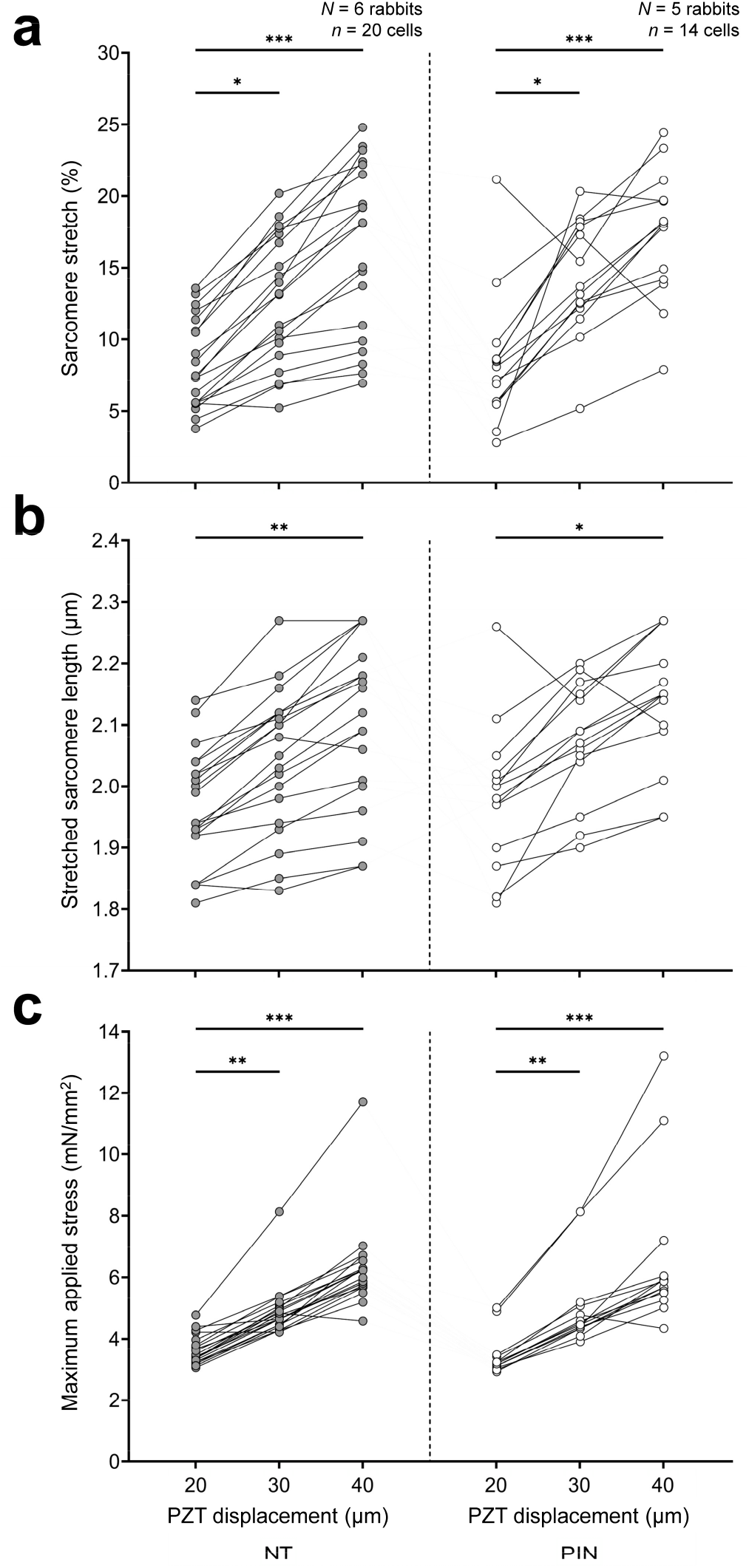
Mechanical characteristics of cell stretch. **a,** Percent sarcomere stretch, **b,** stretched sarcomere length, and **c,** maximum applied stress during acute diastolic stretch of rabbit isolated left ventricular cardiomyocytes exposed to Tyrode (NT, grey, left) or pinacidil (PIN, white, right, 50 µM, to activate ATP-sensitive potassium channels) with increasing levels of piezo-electric translator (PZT) displacement (20, 30, and 40 μm). Differences assessed by one-way ANOVA, with Tukey *post-hoc* tests for **a** and **b**, and by Kruskal-Wallis test for **c**. \**p* < 0.05, \*\**p* < 0.01, \*\*\**p* < 0.001. *N* = rabbits / group, *n* = cells / group.

**Extended Data Figure 2.**
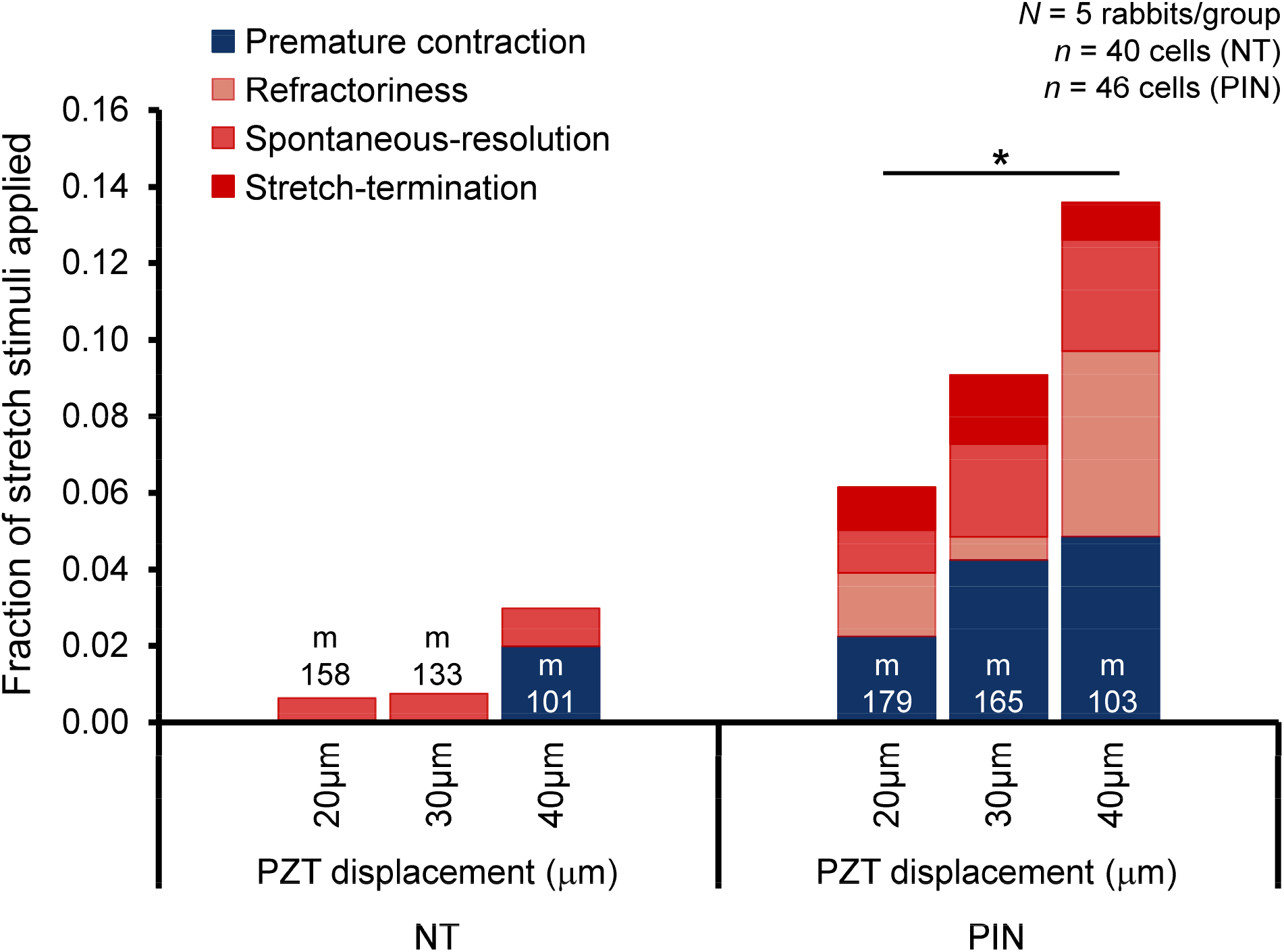
Effect of stretch magnitude on the incidence of mechano-arrhythmogenesis. Incidence of premature contractions (blue) and other self-sustained arrhythmic activity (shades of red) with rapid stretch at increasing levels of piezo-electric translator (PZT) displacement in rabbit isolated left ventricular cardiomyocytes exposed to either Tyrode (NT, left) or pinacidil (PIN, right, 50 µM, to activate ATP-sensitive potassium channels). Differences assessed using chi-square contingency tables and Fisher’s exact test. \**p* < 0.05. *N* = rabbits / group, *n* = cells / group, *m* = stretch stimuli applied.

**Extended Data Figure 3.**
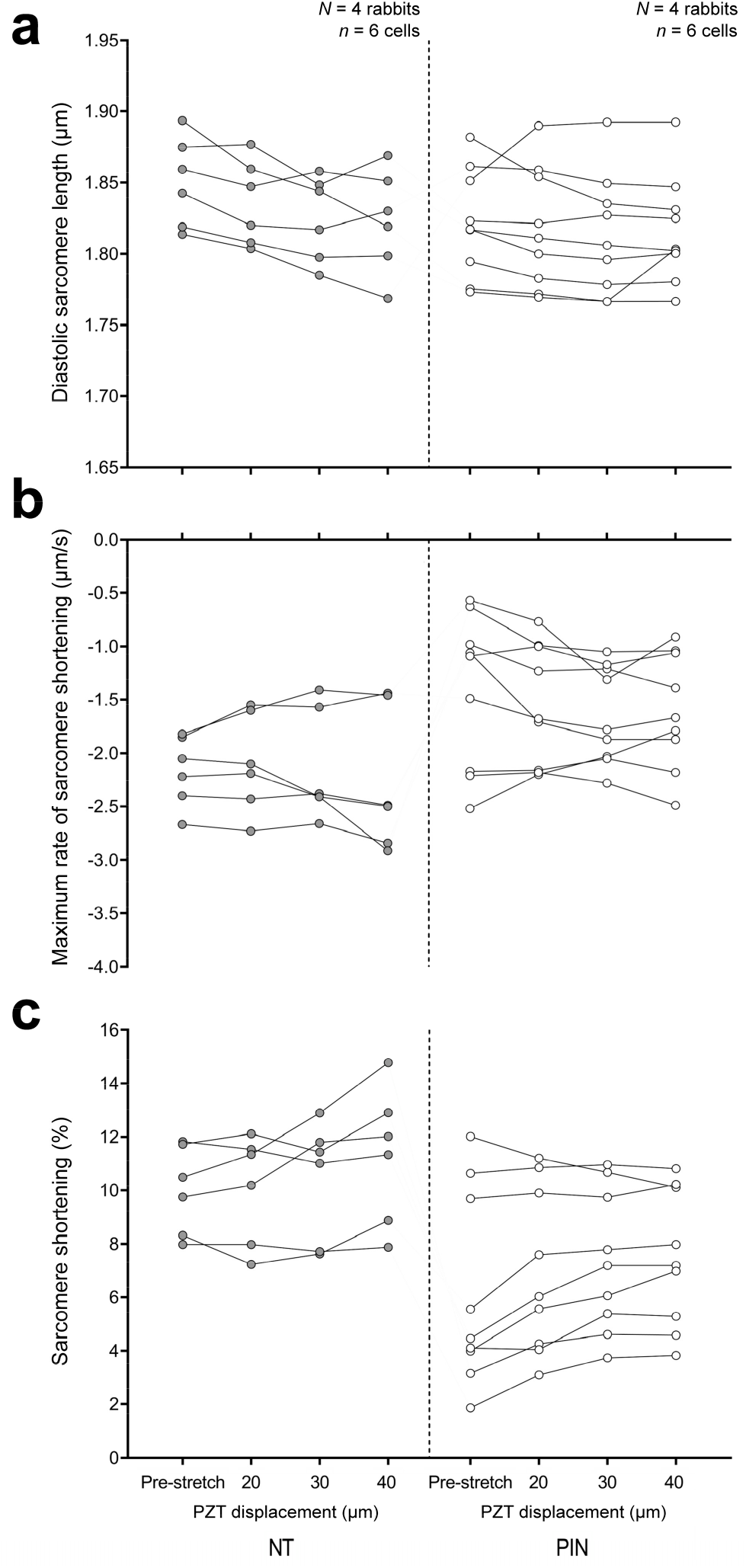
Contractile function before and after stretch in control cells, and in cells with pinacidil (PIN)-induced disruption of repolarisation-relaxation coupling (RRC). **a,** Diastolic sarcomere length and **b,** maximum rate and **c,** percent sarcomere shortening during contraction of rabbit isolated left ventricular cardiomyocytes exposed to either Tyrode (NT, grey, left) or PIN (white, right, 50 µM, to activate ATP-sensitive potassium channels) before (pre-stretch) or after three successive diastolic stretch applications, generated by piezo-electric translator (PZT) displacement (three stretch amplitudes applied were 20, 30 or 40 μm). Measurements for each point averaged over 5 electrically paced contractions. Differences assessed by one-way ANOVA, with Tukey *post-hoc* tests*. N* = rabbits / group, *n* = cells / group.

**Extended Data Figure 4.**
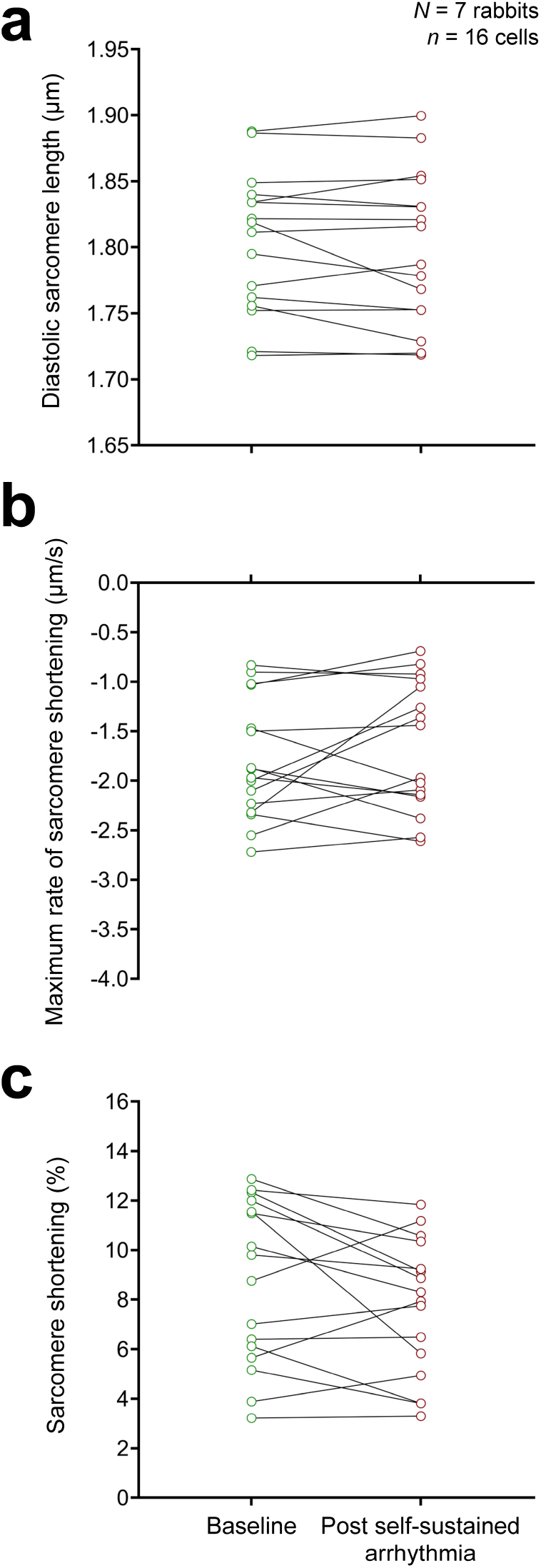
Contractile function after termination of periods with sustained arrhythmic activity. **a,** Diastolic sarcomere length and **b,** maximum rate and **c,** percent sarcomere shortening during contraction of rabbit isolated left ventricular cardiomyocytes in the presence of pinacidil (50 µM, to activate ATP-sensitive potassium channels) before (green) and after (red) termination of stretch-induced self-sustained arrhythmic activity. Measurements for each point averaged over 5 electrically paced contractions. Differences assessed using two-tailed, paired Student’s t-test*. N* = rabbits, *n* = cells.

**Extended Data Figure 5.**
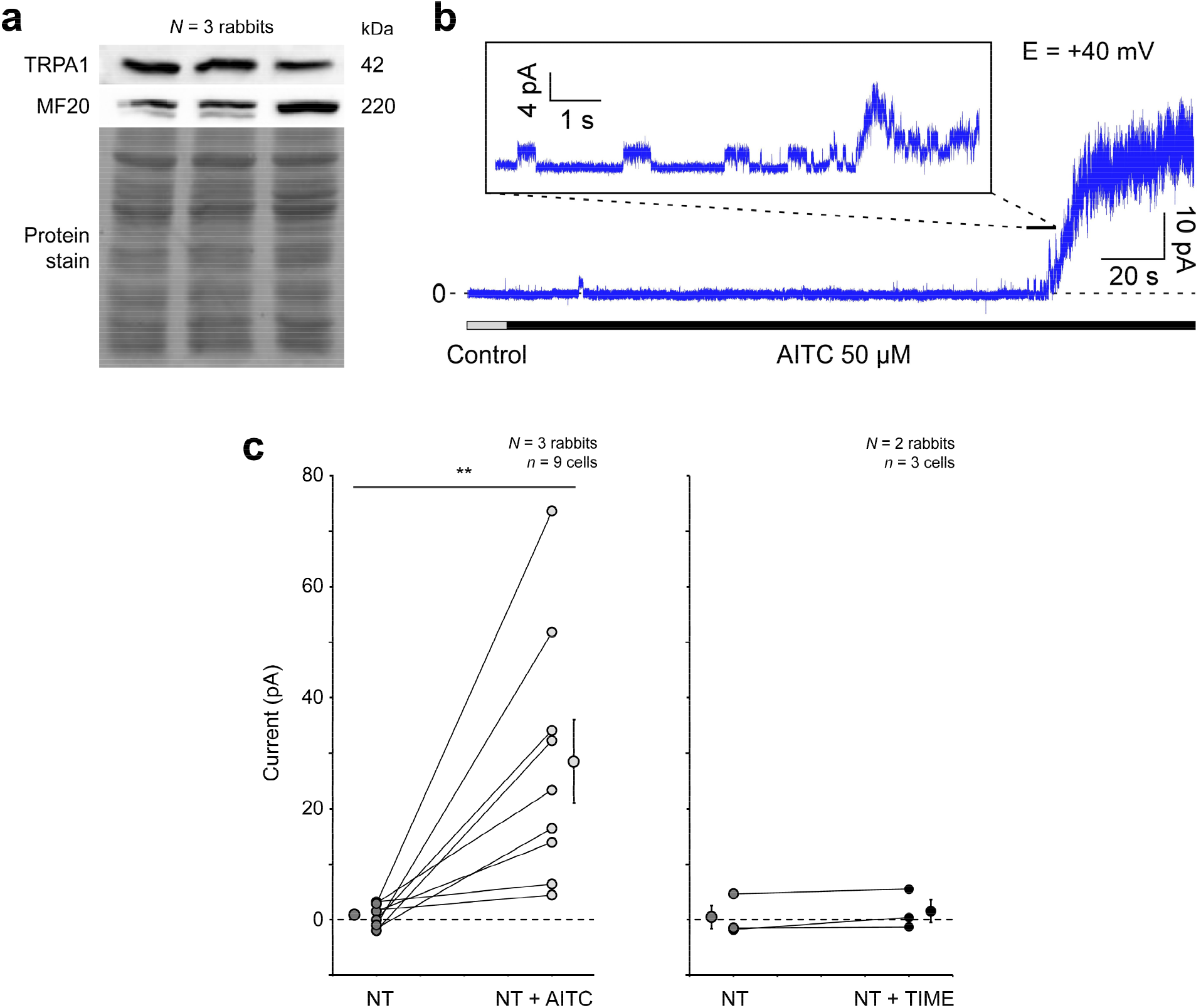
Expression of transient receptor potential ankyrin 1 (TRPA1) channels in rabbit left ventricular myocardium. **a**, Western blot of TRPA1 and myosin heavy chain (MF20) protein expression in rabbit left ventricular free wall tissue. **b**, Representative recording of channel activation in response to allyl isothiocyanate (AITC, 50 μM, to activate TRPA1 channels) in a cell-attached patch (holding potential +40 mV). **Inset**, Detail of channel activation with visible single channel events. **c**, Quantification of AITC-induced current changes (left) and comparison to time-matched same-batch control cells (right). Differences assessed with two-tailed, paired Student’s t-test. \*\**p* < 0.01. Error bars represent standard error of the mean. *N* = rabbits / group, *n* = cells / group.

**Extended Data Figure 6.**
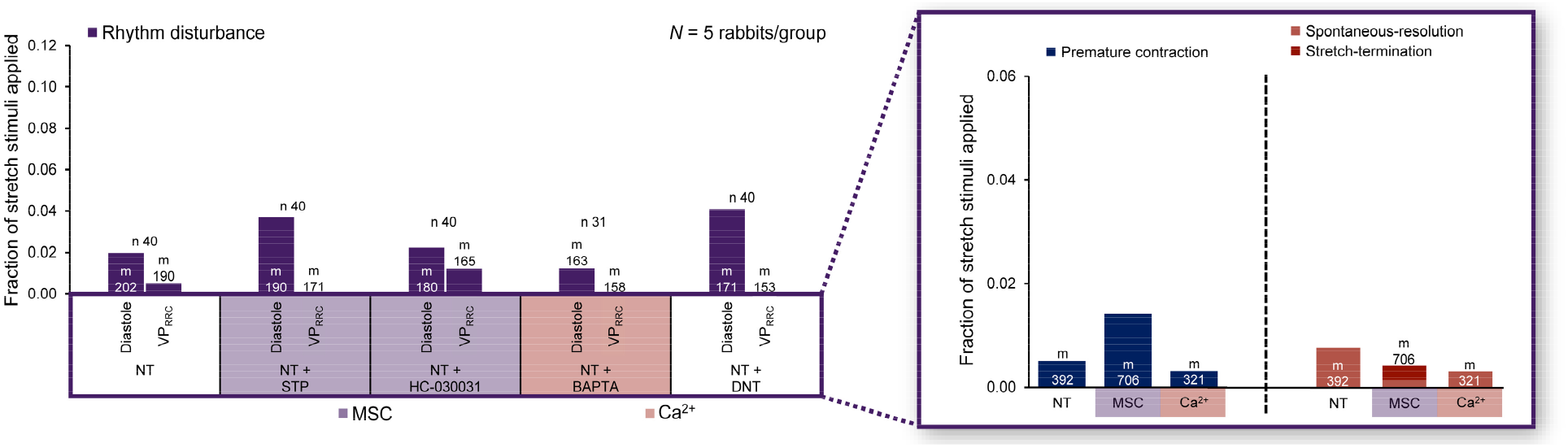
Effect of pharmacological interventions on mechano-arrhythmogenesis with normal repolarisation-relaxation coupling (RRC). Incidence of arrhythmias (purple) in rabbit isolated left ventricular cardiomyocytes without a pinacidil-induced disruption of RRC, following acute stretch applied during diastole or the vulnerable period (VP_RRC_) and exposure to one of the following treatments: Tyrode (NT); NT + streptomycin (STP, 50 µM, to non-specifically block mechano-sensitive channels, MSC); NT + HC-030031 (10 µM, to block transient receptor potential ankyrin 1, TRPA1, channels); NT + BAPTA (1 µM, to buffer cytosolic Ca^2+^, [Ca^2+^]_i_), or NT + dantrolene (DNT, 1 µM, to stabilise ryanodine receptors). **Inset,** Of the total arrhythmia incidence (measured as the sum of diastolic and VP_RRC_ incidence), the proportion of arrhythmias that were premature contractions (blue, left) or self-sustained arrhythmic events (shades of red, right), and the effect of blocking either MSC (measured as the pooled stretches in the NT + STP intervention) or chelating [Ca^2+^]_i_ (measured as the pooled stretches in the NT + BAPTA intervention) on their incidence. Differences assessed using chi-square contingency tables and Fisher’s exact test. *N* = rabbits / group, *n* = cells / group, *m* = stretch stimuli applied.

**Extended Data Figure 7.**
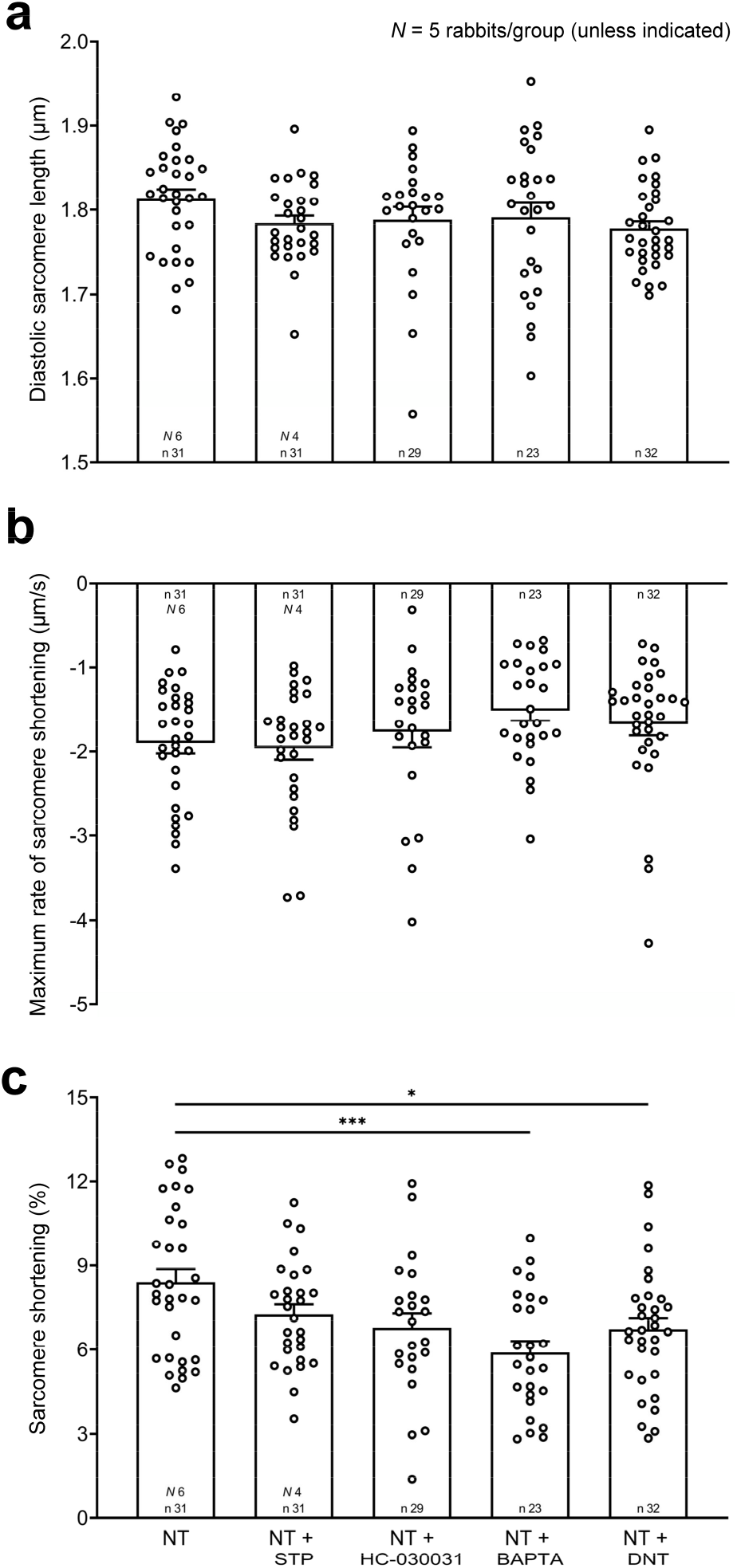
Effect of pharmacological interventions on cellular contractile function in the absence of a pinacidil-induced disturbance of repolarisation-relaxation coupling (RRC). **a**, Diastolic sarcomere length and **b**, maximum rate and **c**, percent sarcomere shortening during contraction of rabbit isolated left ventricular cardiomyocytes exposed to one of the following conditions: Tyrode (NT); NT + streptomycin (STP, 50 µM, to non-specifically block mechano-sensitive channels); NT + HC-030031 (10 µM, to block TRPA1 channels); NT + BAPTA (1 µM, to buffer cytosolic Ca^2+^); or NT + dantrolene (DNT, 1 µM, to stabilise ryanodine receptors). Differences assessed by one-way ANOVA, with Tukey *post-hoc* tests for **a** and **b** and by Kruskal-Wallis test for **c**. \**p* < 0.05, *\*\*\*p* < 0.001. Error bars represent standard error of the mean. *N* = rabbits / group, *n* = cells / group.

**Extended Data Figure 8.**
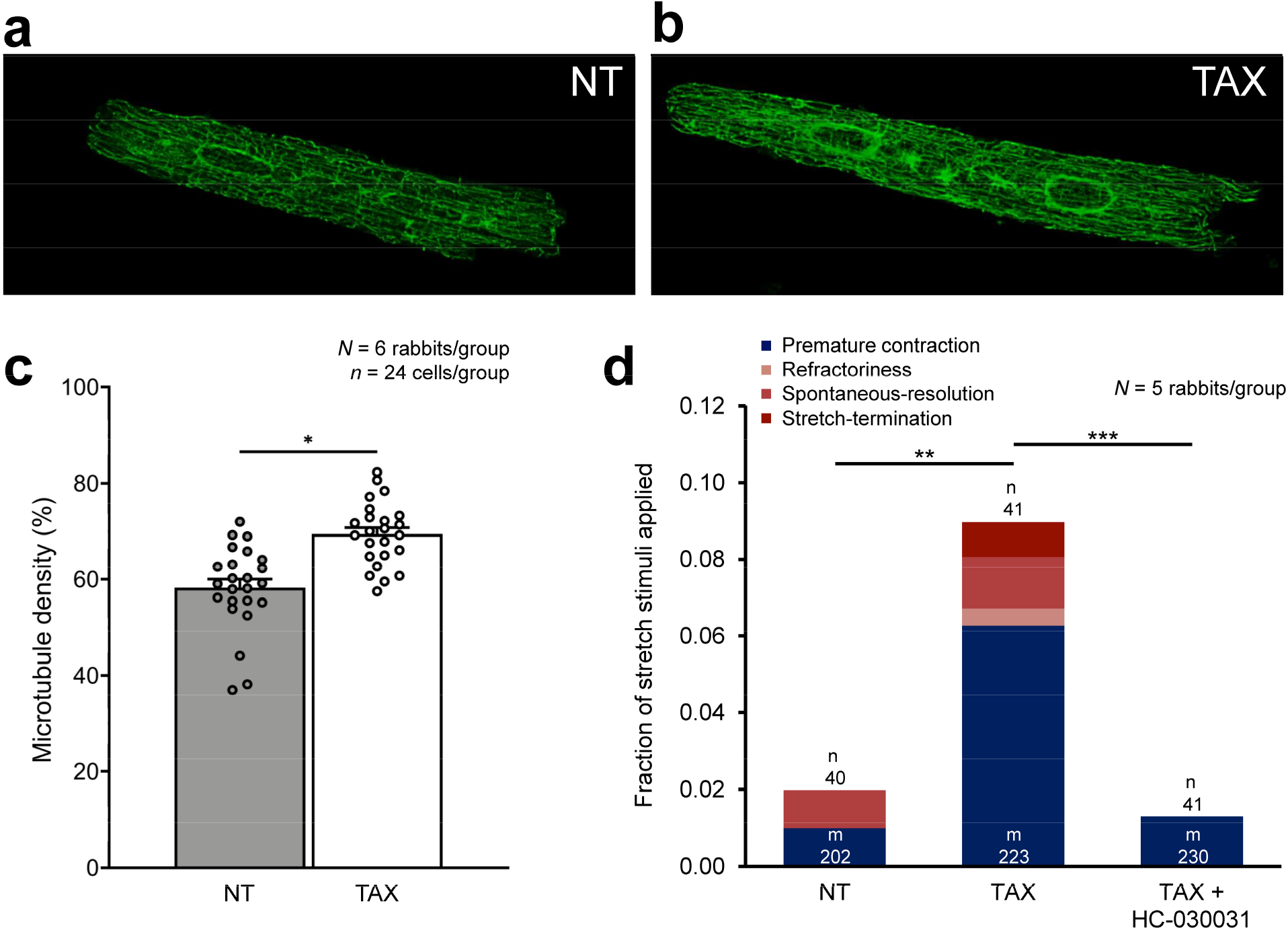
Effect of microtubule density on TRPA1-mediated mechano-arrhythmogenesis. Confocal immunofluorescence images of stained microtubules in rabbit isolated left ventricular cardiomyocytes in either **a,** control conditions (Tyrode, NT) or **b,** following paclitaxel exposure (TAX, 10 µM, to increase the polymerisation and stabilisation of microtubules). **c,** Quantification of microtubule density assessed as the microtubule positive fraction of total cell area. Differences assessed with two-tailed, unpaired Student’s t-test. \**p* < 0.05. Error bars represent standard error of the mean. **d,** Incidence of premature contractions (blue) and other self-sustained arrhythmic activity (shades of red) after acute diastolic stretch of rabbit isolated left ventricular cardiomyocytes exposed to one of the following treatments: NT; TAX; or TAX + HC-030031 (10 µM, to block TRPA1 channels). Differences assessed using chi-square contingency tables and Fisher’s exact test. \**p* < 0.05, \*\**p* < 0.01, \*\*\**p* < 0.001. *N* = rabbits / group; *n* = cells / group, *m* = stretches.

**Extended Data Figure 9.**
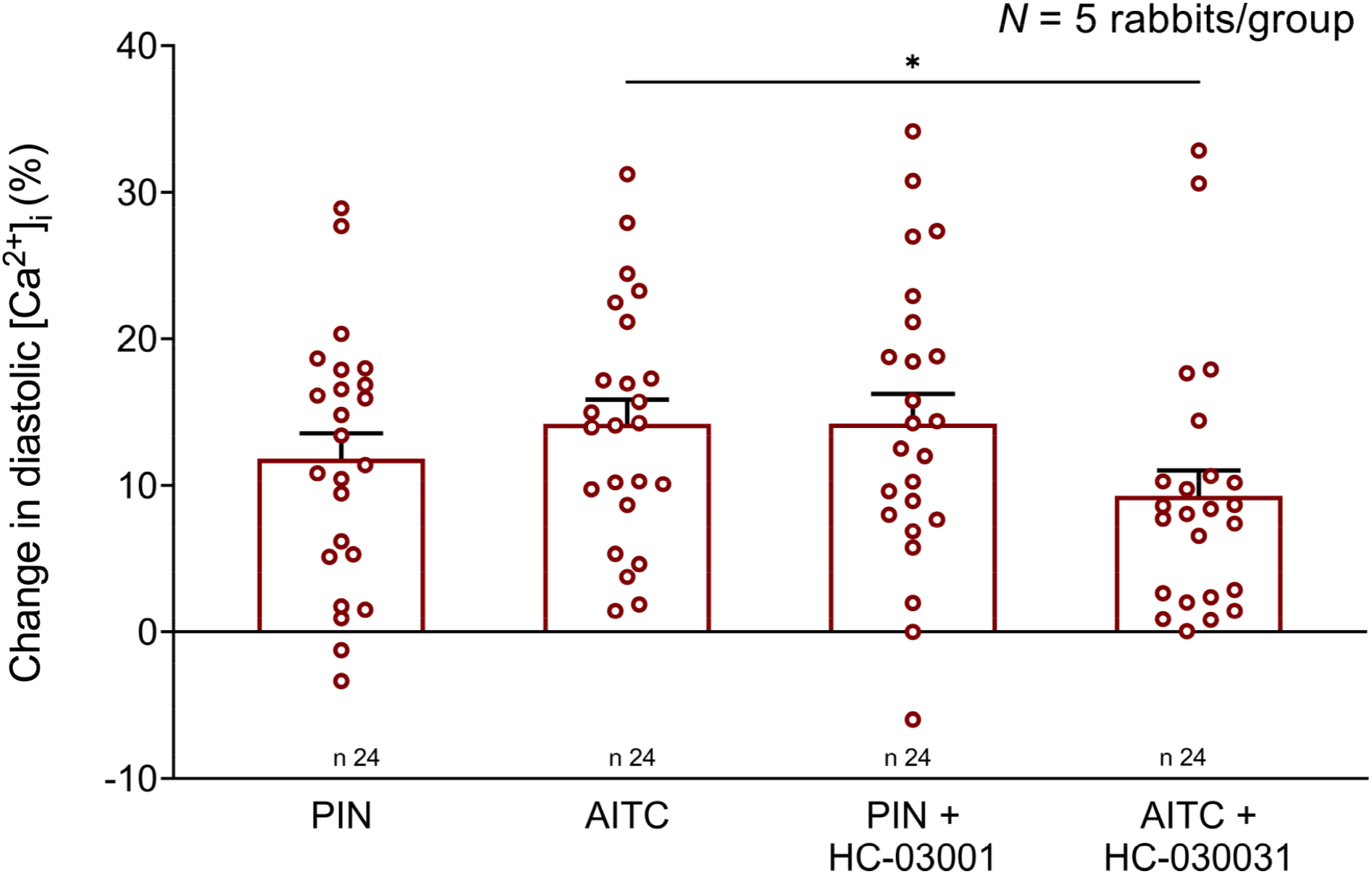
Effect of TRPA1 channel activation on cytosolic Ca^2+^ concentration ([Ca^2+^]_i_). Percent change in diastolic [Ca^2+^]_i_ after exposure to one of the following treatments: pinacidil (PIN, 50 µM, to activate ATP-sensitive potassium channels); allyl isothiocyanate (AITC, 50 μM, to activate TRPA1 channels); PIN + HC-030031 (10 µM, to block TRPA1 channels); or AITC + HC-030031. Differences assessed using two-tailed, unpaired Student’s t-test between groups within PIN or AITC treatment. \**p* < 0.05. Error bars represent standard error of the mean. *N* = rabbits / group, *n* = cells / group.

**Extended Data Figure 10.**
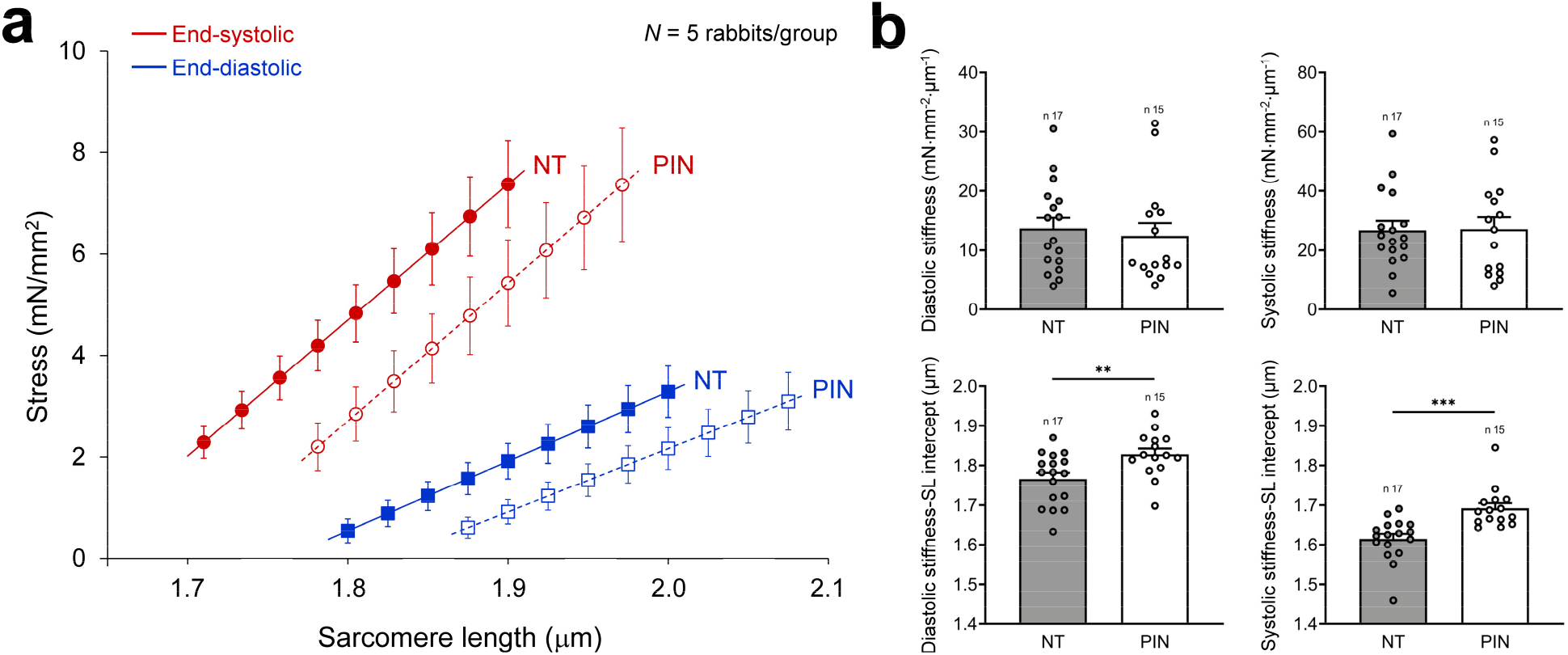
Effect of pinacidil (PIN) on cellular mechanics. **a,** Diastolic (represented by the slope of the end-diastolic stress-length relationship, blue lines with square symbols) and systolic (represented by the slope of the end-systolic stress-length relationship, red lines with round symbols) stiffness of rabbit isolated left ventricular cardiomyocytes either in control conditions (Tyrode, NT; solid lines and filled symbols), or following exposure to PIN (50 µM, to activate ATP-sensitive potassium channels; dashed lines and open symbols). **b,** Calculated diastolic and systolic stiffness, and stiffness-sarcomere length (SL) intercepts in cells exposed to either NT (grey) or PIN (white). Differences assessed using two-tailed, unpaired Student’s t-test. *\*\*p* < 0.01, *\*\*\*p* < 0.001. Error bars represent standard error of the mean. *N* = rabbits / group, *n* = cells / group.

**Extended Data Figure 11.**
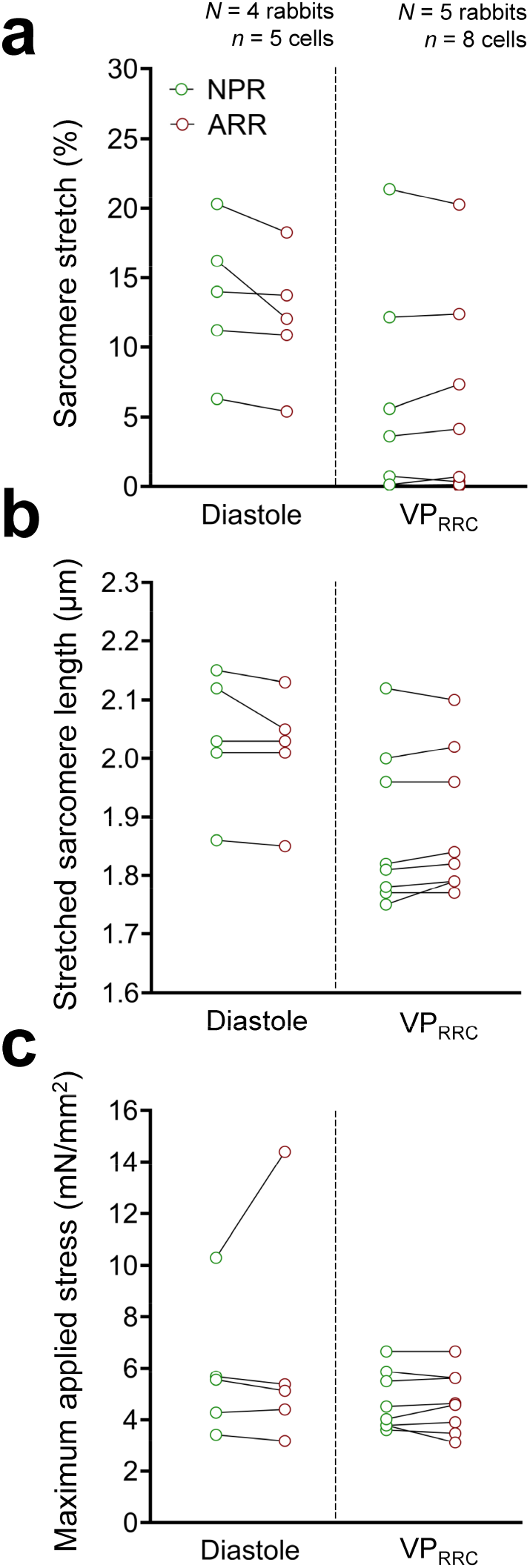
Mechanical characteristics of cell stretch, grouped by timing relative to the electrical cycle and differing electrophysiological outcomes. **a,** Percent sarcomere stretch, **b,** stretched sarcomere length, and **c,** maximum applied stress in pinacidil-treated (PIN, 50 µM, to activate ATP-sensitive potassium channels) cells for two consecutive acute stretch applications either during diastole or the VP_RRC_, where one stretch caused an arrhythmia (ARR, red) and the other did not (cell remained in normal paced rhythm, NPR, green), but not always in that order. Differences assessed using the Wilcoxon matched-pairs test. *N* = rabbits / group, *n* = cells / group.

## SUPPLEMENTARY INFORMATION

**Supplementary Video 1 | Stretch-induced premature contraction.** Example of rapid stretch of a rabbit isolated left ventricular myocyte treated with pinacidil (50 µM, to activate ATP-sensitive potassium channels), applied using microscopic carbon fibres adhered to either end of the cell, which resulted in a premature contraction. Sarcomere dynamics and carbon fibre positions were simultaneously measured to assess arrhythmic incidence and confirm maintenance of contractile function (representative sarcomere trace shown in Fig. 2d).

**Supplementary Video 2 | Stretch-induced refractoriness.** Example of rapid stretch of a rabbit isolated left ventricular myocyte treated with pinacidil (50 µM, to activate ATP-sensitive potassium channels), applied using microscopic carbon fibres adhered to either end of the cell, which resulted in refractoriness. Sarcomere dynamics and carbon fibre positions were simultaneously measured to assess arrhythmic incidence and confirm maintenance of contractile function (representative sarcomere trace shown in Fig. 2f).

**Supplementary Video 3 | Stretch-induced sustained arrythmia with termination by stretch.** Example of rapid stretch of a rabbit isolated left ventricular myocyte treated with pinacidil (50 µM, to activate ATP-sensitive potassium channels), applied using microscopic carbon fibres adhered to either end of the cell, which resulted in a sustained arrhythmia that was terminated by application of an additional stretch. Sarcomere dynamics and carbon fibre positions were simultaneously measured to assess arrhythmic incidence and confirm maintenance of contractile function (representative sarcomere trace shown in Fig. 2h).

**Supplementary Video 4 | Stretch-induced, calcium-driven sustained arrhythmia.** Example of simultaneous voltage (blue)-calcium (red) fluorescence imaging in a rabbit isolated ventricular myocyte treated with pinacidil (50 µM, to activate ATP-sensitive potassium channels). Rapid stretch during the vulnerable period (green bar) resulted in sustained arrhythmic activity, in which oscillations in cytosolic calcium preceded changes in voltage, indicating calcium-driven activity (see inset and Fig. 4a).

## Notes

### Competing Interest Statement

The authors have declared no competing interest.

